# Nanoscale colocalization of NK cell activating and inhibitory receptors controls signal integration

**DOI:** 10.1101/266429

**Authors:** David Tomaz, Pedro Matos Pereira, Nadia Guerra, Julian Dyson, Keith Gould, Ricardo Henriques

## Abstract

NK cell responses depend on the balance of signals from inhibitory and activating receptors. However, how the integration of antagonistic signals occurs upon NK cell-target cell interaction is not fully understood. Here, we provide evidence that NK cell inhibition via the inhibitory receptor Ly49A is dependent on its relative colocalization at the nanometer scale with the activating receptor NKG2D upon immune synapse (IS) formation. NKG2D and Ly49A signal integration and colocalization was studied using NKG2D-GFP and Ly49A-RFP-expressing primary NK cells, forming ISs with NIH3T3 target cells, with or without expression of single chain trimer (SCT) H2-D^d^ and an extended form of SCT H2-D^d^-CD4 MHC-I molecules. Nanoscale colocalization was assessed by Förster resonance energy transfer (FRET) between NKG2D-GFP and Ly49A-RFP and measured for each synapse. In the presence of their respective cognate ligands, NKG2D and Ly49A colocalize at a nanometer scale leading to NK cell inhibition. However, increasing the size of the Ly49A ligand reduced the nanoscale colocalization with NKG2D consequently impairing Ly49A-mediated inhibition. Thus, our data shows NK cell signal integration is critically dependent on the dimensions of NK cell ligand-receptor pairs by affecting their relative nanometer-scale colocalization at the IS. Together, our results suggest the balance of NK cell signals, and NK cell responses, are determined by the relative nanoscale colocalization of activating and inhibitory receptors in the immune synapse.

## Introduction

NK cell activation is dependent on a different array of germline-encoded receptors capable of triggering effector responses against infection (1–3) and cellular transformation (4, 5), and yet maintaining tolerance towards healthy cells and tissues (6, 7). NK cell functionality is widely characterized by a balance between activating and inhibitory receptors (8, 9). However, upon immune synapse formation, how NK cells integrate signals from functionally antagonistic receptors, is not yet fully understood (10). One major hypothesis to explain immune signal integration is based on the kinetic segregation (K-S) model (11). This model states that the balance of antagonistic signals is mediated by the equilibrium of phosphorylation (kinases) and dephosphorylation (phosphatases), which depends on the relative colocalization of receptors and their associated signaling molecules upon synapse formation. This hypothesis has been validated using T cells, in the context of TCR triggering (12, 13) and, more recently, NK cells (14, 15). The K-S model suggests immune inhibition, upon immune synapse formation, is maintained by the dephosphorylation of tyrosines in stimulatory receptors (e.g ITAMs-associated receptors such as TCRCD3 or NKG2D-DAP12), due to a nanoscale colocalization with receptors bearing phosphatase activity (e.g. CD45, CD148) or with the capacity of recruiting phosphatases such as SHP-1/2 or SHIP (e.g. ITIM-associated receptors such as PD1, KIR2DL2 or Ly49A) (16, 17). Several recent studies have investigated the relationship between immune signal integration and colocalization of antagonistic signal receptors. Using altered dimensions of receptor-ligand pairs it has been shown that the molecular dimensions of receptorligand pairs have a proportional, size-dependent effect on immune cell signal integration and functionality in both T and NK cells. In T cells, TCR-MHC-I dimensions have been shown to play a major role in triggering TCR activation (12). The molecular dimensions of CD45 and CD148 have also been reported to affect T cell functionality (18). More recently, the initiation of T cell signaling was confirmed by the CD45 segregation at close contact points (19). In NK cells, elongation of activating and inhibitory NK cell receptor ligands has been shown to affect NK cell activation and inhibition, respectively (14, 15). These studies showed elongation of H60a reduces NKG2D-mediated activation, without reducing binding. Similarly, elongating ligands for both the inhibitory Ly49C/I and CD94/NKG2A receptors reduced NK cell inhibition (14). In another study using human NK cells, it was found that elongated versions of HLA-Cw6 and MICA reduced their inhibition and activation, respectively (15). This study also suggested that a relative colocalization of matched-size NKG2D/MICA and KIR2DL1/HLA-Cw6 complexes is required for efficient NK cell signal integration. Although multiple combinations of NK cell activating and inhibitory receptors have been identified (20–22), their relative contributions to NK cell responses need to be further investigated. A murine model study by Regunathan *et al* (2015), investigated the balance of signals between Ly49A and NKG2D. This study showed that NKG2D-dependent activation could be counterbalanced by inhibitory receptors, in particular, Ly49A (23). The interplay of signaling strength between the inhibitory receptors, Ly49A/G, and the activating receptor, NKG2D, was shown to determine the magnitude of cell activation. Ly49A was effective at inhibiting NKG2D-dependent responses, even in the presence of high levels of NKG2D ligand expression (23). Ly49A-mediated inhibition is hypothesized to occur due to its relative proximity to activating receptors (e.g. NKG2D), and the proximal recruitment of Src homology region 2 domain-containing phosphatase-1 and-2 (SHP1/2) or phosphatidylinositol-3,4,5-trisphosphate 5-phosphatase 1 (SHIP) (24, 25). However, a nanometer scale colocalization of Ly49A with activating receptors, upon immune synapse formation, in primary NK cells, has not yet been reported. In the present study, we used retrogenic mice to generate primary NK cells expressing fluorescent protein (FP)-tagged Ly49A and NKG2D receptors. The colocalization of NKG2D-GFP and Ly49A-RFP was investigated on a nanometer scale by measuring Förster resonance energy transfer (FRET) between the FP-tagged receptors upon synapse formation. Interestingly, we observed colocalization of Ly49A and NKG2D upon IS formation, these results suggest the balance of NK cell signals might be determined by the relative colocalization of activating and inhibitory receptors at a nanometer scale. These new findings are important as they suggest a molecular mechanism of balance of NK cell signals dependent on a differential organization of receptors upon synapse formation.

## Results

### Generation of NK cells co-expressing NKG2D-GFP and Ly49A-RFP *in vivo*

In this work, we created retrogenic (RT) mice displaying NK cells expressing NKG2D-GFP and Ly49A-RFP receptors. Several previous studies have used FP-tagged NK cell receptors or ligands to study NK cell signaling. These studies expressed NKG2D- and Ly49A-tagged receptors on T and NK-like cell lines (26, 27). In this study, we created authentic primary C57BL/6 (B6) NK cells expressing FP-tagged NK cell receptors *in vivo*. The RT mice technique has previously been used to create specific TCR- and BCR-expressing T and B cells, respectively (28, 29), but here we applied it for the first time to create NK cells expressing FP-tagged receptors *in vivo* (Supplementary Figure S1). pMIGR1 expressing the long and short isoforms of NKG2D-GFP (NKG2D-L-GFP and NKG2D-S-GFP), and Ly49A-RFP were generated (Figure 1a-b). By retroviral transduction, we expressed NKG2D-GFP and Ly49A-RFP in three different immune cells: i) B3Z CD8+ T cell hybridoma, (ii) Hematopoietic progenitor cells (HPC)-derived NK cells, and (iii) NK cells *in vivo*, from RT mice (Figure 1c). NK cells from RT mice were generated as previously described (30, 31). 293 T cells were efficiently transfected with NKG2D-GFP and Ly49A-RFP (Supplementary Figures S2 and S3). HPCs from NKG2D-deficient mice were transduced and used as donor cells (Supplementary Figures S4 and S5). NKG2D-deficient donor cells were used to avoid the expression of endogenous untagged NKG2D. HPCs were first transduced and then differentiated *in vitro* into HPC-derived NK cells as previously described. B3Z CD8+ T cell hybridoma cells were directly transduced. Both NKG2D-GFP and Ly49A-RFP were expressed at the cell surface (Figures 2 a-b), and RFP and GFP intensities were correlated with cell surface expression levels of Ly49A and NKG2D, respectively (Figure 1c). The cell surface expression of NKG2D isoforms varied between CD8+ T cells and NK cells, where we observed the relative percentage of NKG2D-expressing cells was higher on NK cells than CD8+ T cell hybridomas. NKG2D-S-GFP presented higher levels of expression when compared with NKG2D-L-GFP (Supplementary Figure S6). This is consistent with results obtained for NKG2D in previous studies (32–34). We successfully generated NK cells expressing FP-tagged receptors in B6 mice using the RT mice technique (Figures 1 and 2). NK cells expressing NKG2D-GFP and Ly49A-RFP were detected in murine peripheral blood and spleen for up to four months (Supplementary Figure S7). It is therefore apparent that authentic primary NK cells expressing FP-tagged NK cell receptors can be generated using the RT mice technique. This establishes proof-of-principle for applying this technique to produce NK cells expressing chimeric receptors. Our results also indicate that FP-tagging of NKG2D and Ly49A does not significantly impair their cell surface expression, providing compelling evidence for the application of this technique in colocalization studies.

**Fig. 1.**
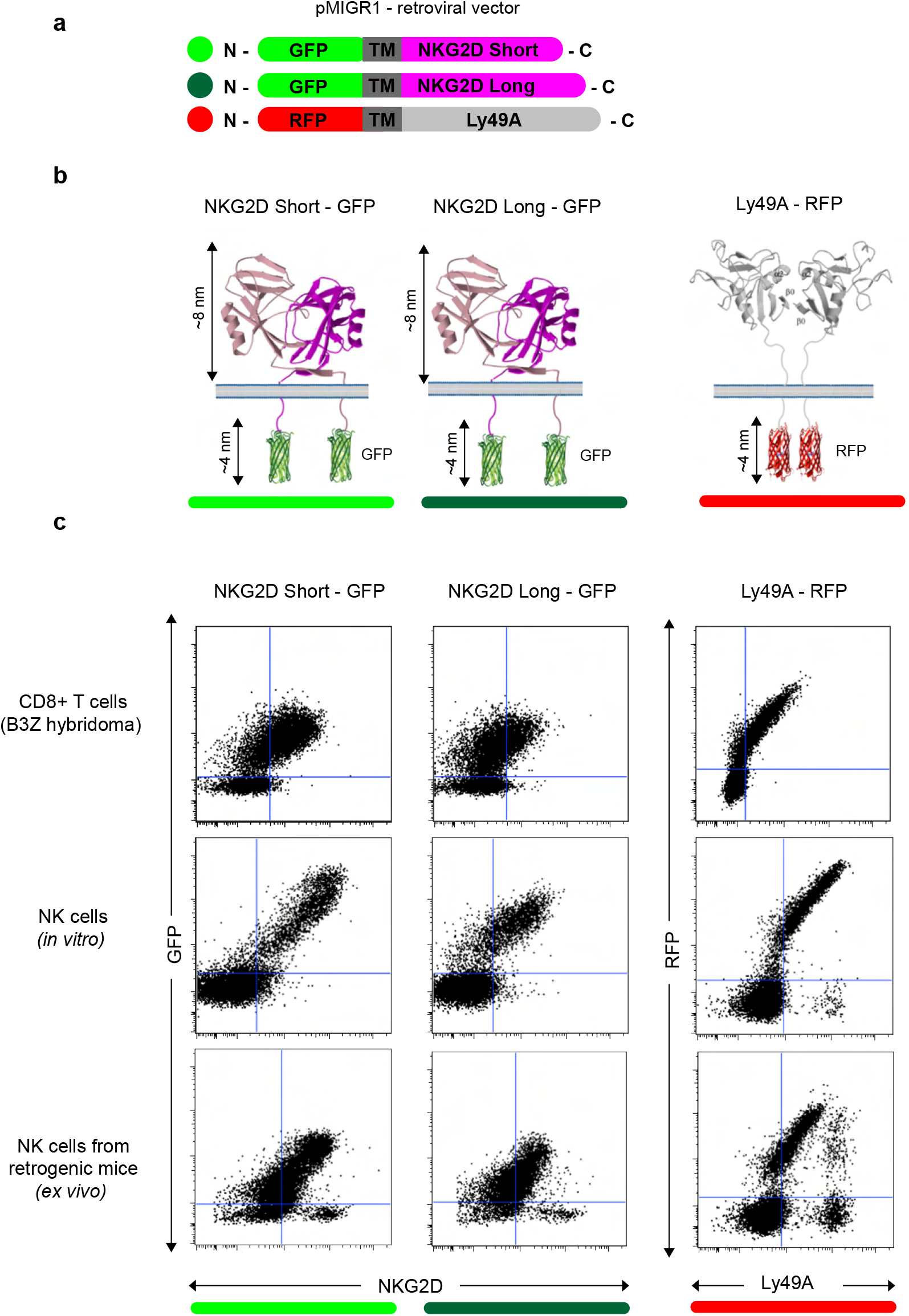
Generation and expression of NKG2D(S/L)-GFP and Ly49A-RFP expressing NK cells *in vivo*. **a**, Schematic representation of the constructs of mouse NKG2D-GFP and Ly49A-RFP used for functional and colocalization studies. All constructs were subcloned into the pMIGR1 retroviral expression vector. **b**, Schematic of NKG2D-GFP and Ly49A-RFP at the NK cell surface. Both molecules form homodimers. Image representation is not to scale. **c**, NK cells expressing NKG2DGFP and Ly49A-RFP generated *in vivo*. Retroviral-mediated expression of NKG2D-GFP (S/L) and Ly49A in murine B3Z CD8+ T cell line (top), NK cells differentiated *in vitro* from HPCs (middle) and NK cells obtained from retrogenic (RT) mice (bottom, see also Supplementary Figure S1). Experiments with NK cells differentiated *in vitro*, were performed by transducing HPCs from NKG2D-deficient C57BL/6 mice (Supplementary Figure S2) and data shown are gated on NK1.1+, NKp46+ and CD3-cells. Splenic NK cells from RT mice were gated as CD3-, CD8-, NK1.1+ and CD49+ (DX5) cells. The data are representative of more than three experiments.

**Fig. 2.**
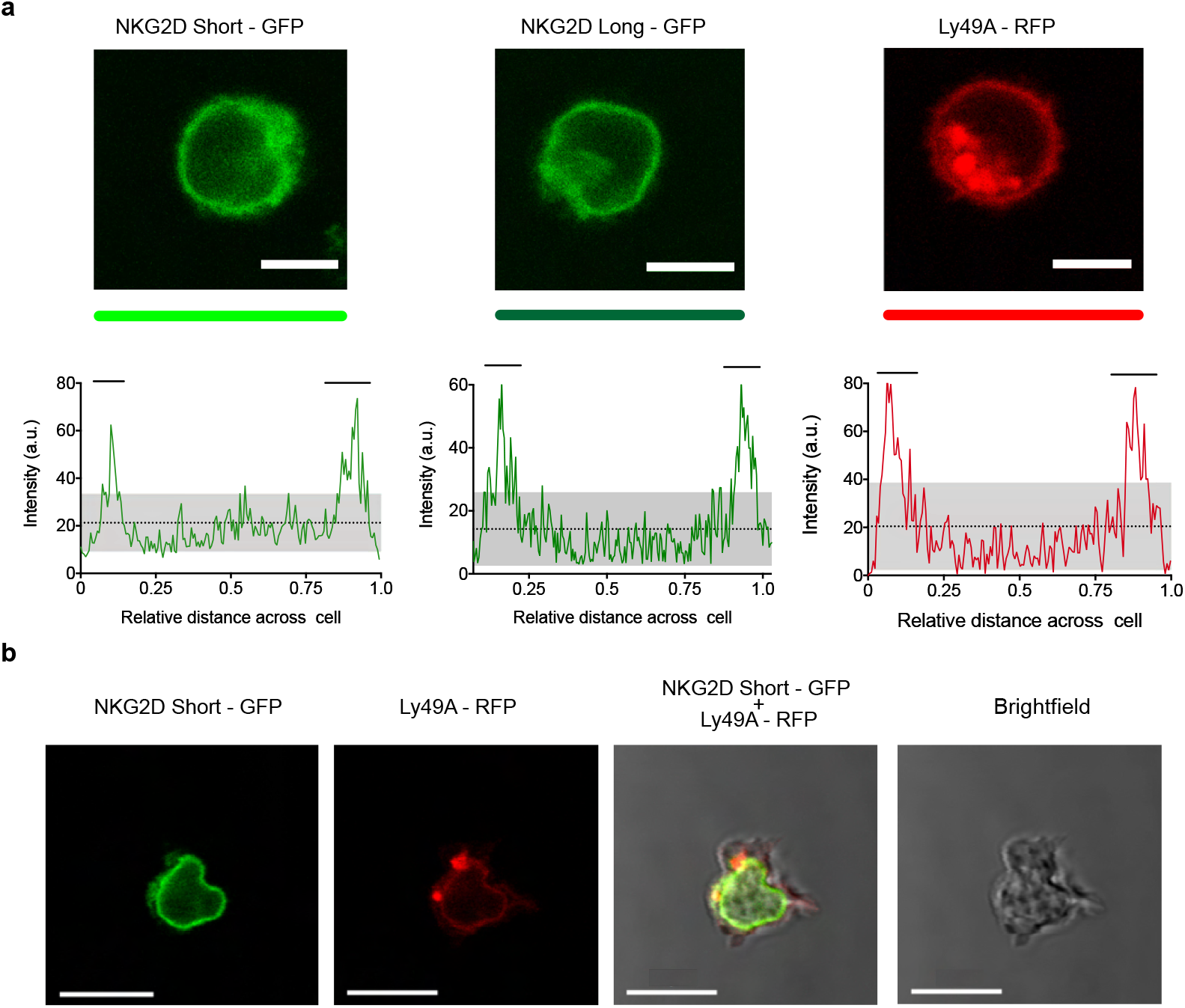
Primary NK cells express NKG2D-GFP and Ly49A-RFP receptors at the cell surface. **a**, Confocal z-stack live cell imaging of NKG2D short-GFP, NKG2D long-GFP and Ly49A-RFP splenic NK cells obtained from retrogenic mice (top). Plot profiles show GFP and RFP signal intensities across cell diameter (bottom). The mean signal intensity is represented by a dotted line. Grey areas represent lower and upper 95% CI of mean. Horizontal bars indicate areas of the plot where signal is above the upper 95% CI of mean. One representative experiment is shown. Scale bars, 5um. **b**, Confocal z-stack live cell imaging of double transduced - NKG2D short-GFP + Ly49A-RFP NK cell (*in vivo*). Scale bars, 10um.

### NKG2D-GFP associates with adaptor proteins DAP10 and DAP12

It is known that the NKG2D-S isoform associates with both DAP10 and DAP12, whereas the NKG2DL isoform associates only with DAP10. DAP10 associated NKG2D isoforms are present in both NK and activated CD8+T cells. DAP12 association with the NKG2D short isoform occurs in NK cells only (32–34). Here we tested the effect of GFP tagging on both NKG2D isoforms and their association with DAP10 and 12. Different NKG2D-GFP isoforms were found to associate with either DAP10 or DAP12 (Figure 3a). The relative percentage of NKG2D+/GFP+ cells was analyzed (Figure 3b). NKG2D-L-GFP was found to associate with DAP10 (P<0.0001) but not with DAP12 (P=0.6232), whereas NKG2D-S-GFP was found to associate with both, but more efficiently with DAP10 (P<0.0001) than DAP12 (P=0.0009). Thus, NKG2D-S-GFP was generally found to associate more efficiently with both adaptor proteins than NKG2D-L-GFP. These differences in adaptor association are consistent with previous studies showing that the cell surface expression of NKG2D requires stabilization by DAP10 or DAP12 molecules (32). However, it is possible that the tagging of GFP onto NKG2D-S may affect its association with DAP12. This would be consistent with the hypothesis that elongating the NKG2D cytoplasmic tail affects DAP12 association, as suggested previously [40]. Thus, it seems that DAP12 association is more dependent on the length of the NKG2D cytoplasmic tail than DAP10. NKG2D-GFP signaling and functionality was not directly tested. Nevertheless, NKG2D-GFP association with DAP10 and DAP12 indicates likely maintenance of functionality. Functionality has also been previously reported for human NKG2D-L fused to GFP (26).

**Fig. 3.**
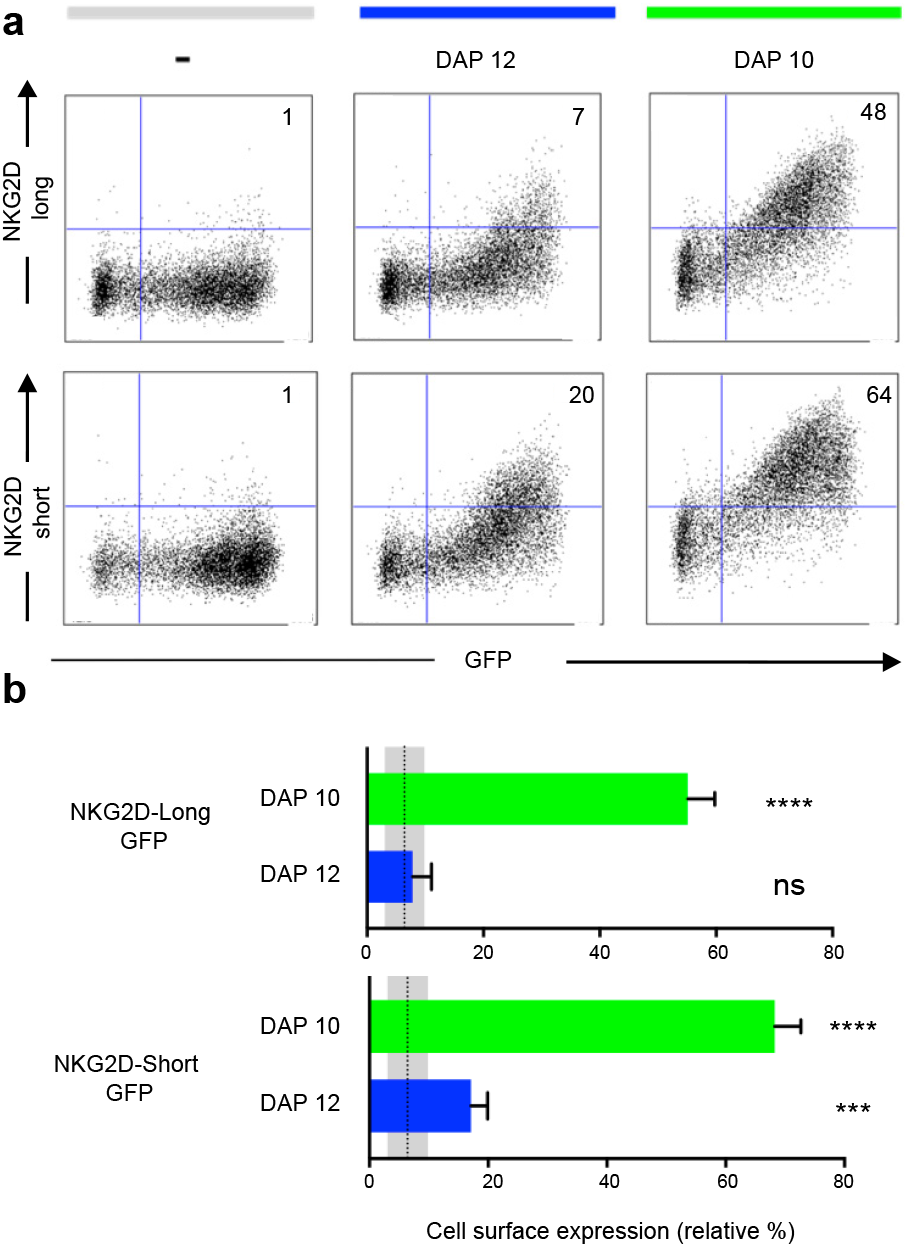
NKG2D-GFP associates with adaptor proteins DAP10 and DAP12. **a**, Expression of NKG2D-GFP short and long isoforms at the cell surface depends on association with DAP10 or DAP12. HEK 293T cells were transiently co-transfected with pTagGFP-NKG2D short or long isoform and either DAP12 or DAP10 as indicated. Cells were analysed by flow cytometry for NKG2D surface expression. Numbers indicate the percentage of cells in the respective quadrant. One representative experiment is shown. **b**, NKG2D-GFP short isoform associates with both DAP10 and DAP12. Both isoforms can associate with DAP10, but DAP12 associates significantly better with the short than with the long NKG2D-GFP isoform. The bar graphs show the mean percentage ± SD from four replicates, and bars with statistically significant difference are as shown: *P<0.05, **P<0.01, ***P<0.001, ****P<0.0001. ns, no statistically significant difference (p>0.05). Unpaired T tests were performed to assess statistical significance in comparison with the untransfected control. The mean is represented by a dotted line and grey areas represent mean percentage ± SD of untransfected control (-).The data are representative of three separate experiments.

### Activating and inhibitory ligand dimensions affect NK cell responses

Earlier studies showed elongated NK cell ligands affect NK cell responses, with elongation of activating and inhibitory ligands affecting NK cell activation and inhibition, respectively (14). In this study, two NK cell ligands were used to test whether activating and inhibitory ligand dimensions affect NK cell responses. Activating, H60a-CD4, and inhibitory, SCT D^d^-CD4, elongated ligands were generated by insertion of four inert human IgSF CD4 domains. Here we demonstrate that elongating H60a weakens NK cell activation and elongating D^d^ reduces NK cell inhibition. These results are consistent with previous studies (14, 15). RMA target cells were transfected with H60a and H60a-CD4 (Fig.4a). Both constructs were found to be expressed on RMA cells with similar cell surface expression (Fig. 4b). Elongation of H60a was observed to reduce NK cell lysis at different effector:target (E:T) cell ratios (5:1, P=0.002; 10:1, P=0.005; 20:1, P<0.0001; Figure 4c). These results were found to be NKG2D-dependent (data not shown). Thus, it is evident in our study that elongated activating NK cell ligands have a detrimental effect on NK cell activation. Next, we investigated whether the elongation of H60a interfered with NKG2D binding and cell-cell conjugate formation. No accumulation of NKG2D at the immune synapse was observed with untransfected RMA cells (Figure 4d). NKG2D accumulation was found to increase at the immune synapse formed by RMA cells expressing either H60a (P=0.0148) or H60a-CD4 (P=0.0162; Figures 4d and 4e). A similar observation was made for cell-cell conjugation; NK cells were found not to form conjugates efficiently with untransfected RMA cells, but they formed conjugates with RMA cells expressing H60a (P<0.001) or elongated H60a-CD4 (P<0.001; Figure 4f). Our results confirm elongation of H60a affects NK cell activation but not by impairing NKG2D binding and cell-cell conjugation. Next, we investigated whether ligand dimensions play a role in NK cell inhibition. Figure 5 represents the impact of the elongation of SCT D^d^ in NK cell inhibition. NIH3T3 cells transfected with SCT Dd and SCT D^d^-CD4 ligands were used as target cells (Fig. 5a). Both constructs were expressed on NIH3T3 cells at similar levels (Fig. 5b, left). SCT D^d^ and SCT D^d^-CD4 molecules were also efficiently transfected and correctly folded on CHO cell surface as demonstrated by antibody staining with two different anti H2-D^d^ monoclonal antibodies (Supplementary Figure S8). NIH3T3 cells were observed to endogenously express the NKG2D ligand, H60a (Figure 5b, right). NIH3T3 expressing SCT D^d^ were more resistant to NK cell lysis by Ly49A+ enriched NK cells at different E:T ratios (1:1, P=0.007; 5:1, P=0.003; 10:1, P=0.0002; Figure 5c). However, the elongation of the Ly49A ligand SCT D^d^ was found to reduce Ly49A-mediated inhibition. Using the elongated SCT D^d^ ligand, lysis was higher and equivalent to the levels of the NIH3T3 untransfected control. These results were in agreement with a loss of function of the elongated ligand also observed for NK cell activation when using H60a-CD4 ligand (see Figure 4). Thus, it is evident that inhibitory NK cell ligand dimensions control NK cell responses in agreement with previous observations (14, 15). The differences in NK cell inhibition are unlikely to be due to different ligand expression since levels of ligand expression were similar (Fig. 5b). It has been reported that elongated ligands can increase the inter-membrane space in the IS (12). Thus, it was hypothesized that ligand elongation affects the nanoscale colocalization of receptors.

**Fig. 4.**
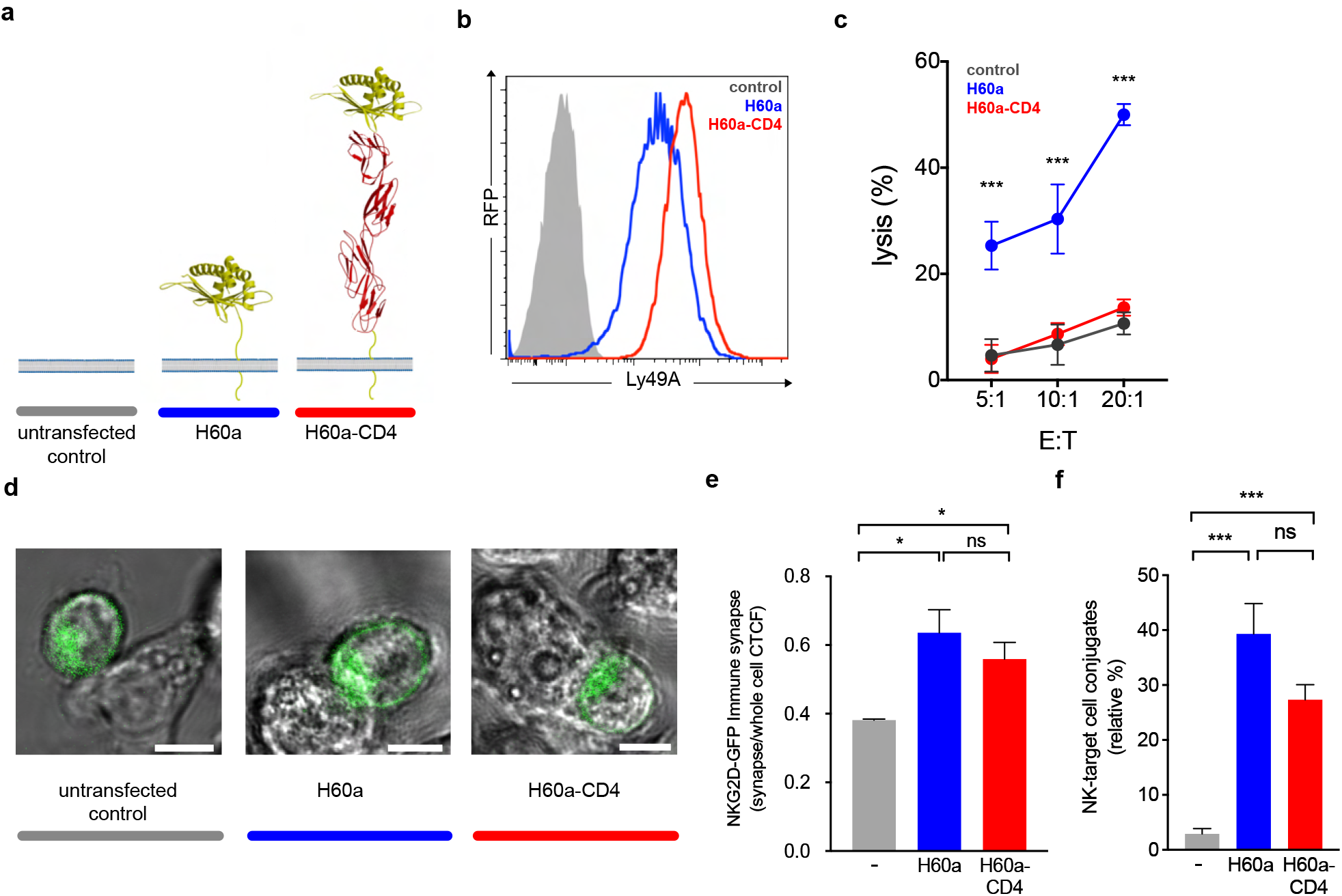
Activating ligand dimensions affect NK cell activation. **a**, Schematic representation of H60a NKG2D ligand and its elongated form. The insertion of four IgSF domains creates a predicted elongation of 12nm. For illustrative purposes the RAE-1B structure was used to represent H60a. **b**, H60 cell surface staining of RMA transfectants (antibody clone 205326, R&D systems). The control (solid grey) corresponds to untransfected RMA cells, H60a (blue) and H60a-CD4 (red) correspond to RMA clones with similar levels of ligand expression. **c**, Elongation of H60a reduces NK cell activation. Untransfected RMA (control, grey), RMA H60a (H60a, blue) and RMA H60a-CD4 (H60a CD4, red) transfected cells expressing similar levels of ligands were used as target cells. IL-2-expanded NK cells from C57BL/6 mice were used in a killing assay at the indicated effector:target (E:T) ratios. Data show mean+SD %lysis (n=3), and groups with statistically significant difference are as shown: *P<0.05, **P<0.01, ***P<0.001, ns, no statistically significant difference (p>0.05). Multiple unpaired t tests were used applying the Holm-Sidak method. Data are representative of two independent experiments. **d**, Confocal z-stack live cell imaging of NKG2D short-GFP NK cells *ex vivo* targeting untransfected, H60a or H60a-CD4-expressing RMA cells. **e**, Elongated H60a supports efficient target cell conjugate formation with NK cells and NKG2D-GFP accumulation at the immune synapse. **f**, NK-target cell conjugates are formed between NK cells and H60a and H60a-CD4 target cells with no significant difference between both these target cells. The data are representative of 3 separate experiments. Untransfected RMA cells (grey); H60a-expressing RMA cells (blue); H60a-CD4-expressing RMA cells (red). CTCF, corrected total cell fluorescence

**Fig. 5.**
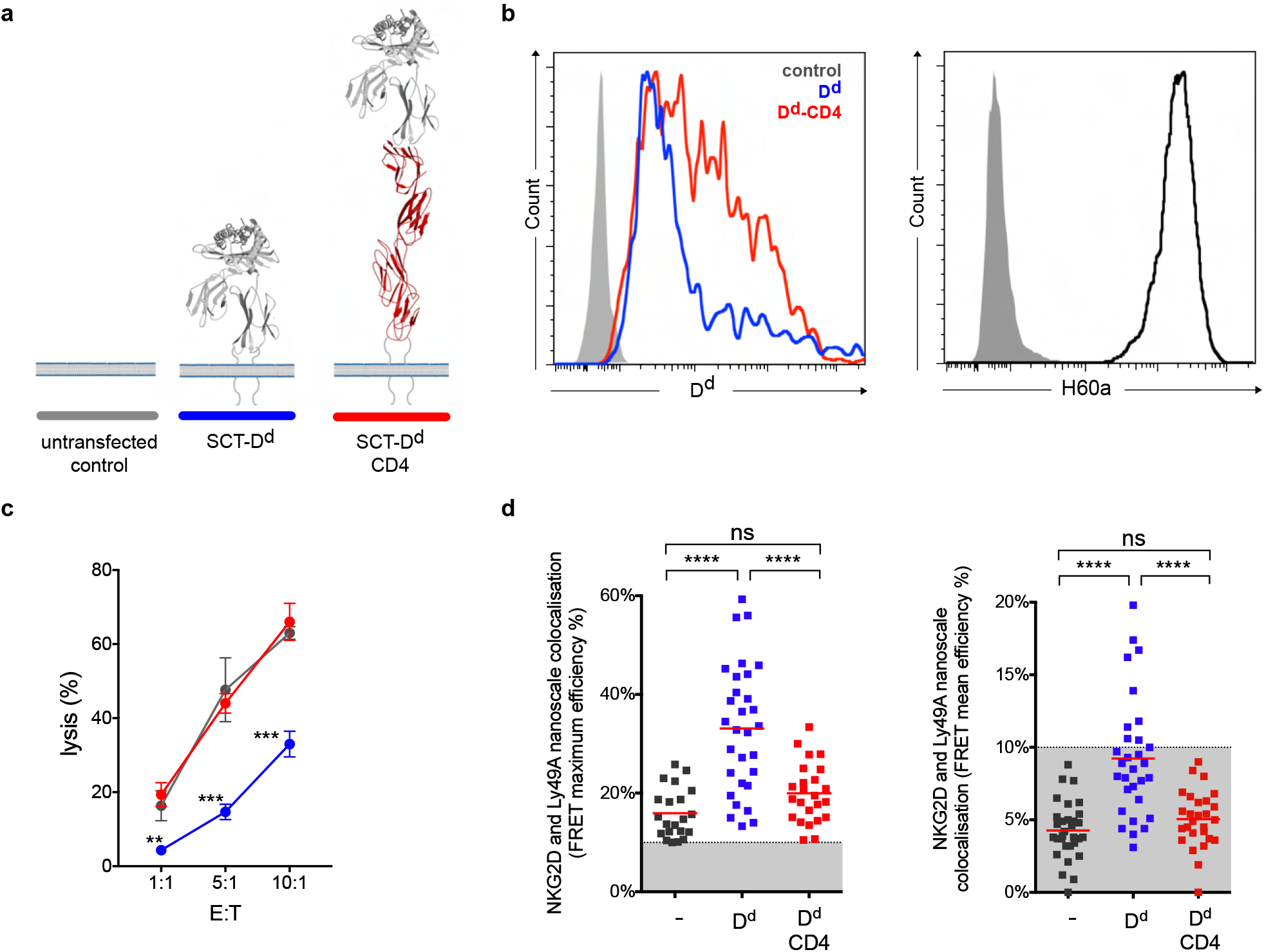
Ly49A-mediated inhibition of NKG2D-mediated NK cell activation correlates with the colocalization of Ly49A with NKG2D at a nanometer-scale. **a**, Schematic representation of SCT D^d^ Ly49A ligand and its elongated form. **b**, H-2-D^d^ cell surface staining (antibody clone 34-2-12) of NIH3T3 transfectants (left panel). The control (solid grey) corresponds to untransfected NIH3T3 cells, D^d^ (blue) and D^d^-CD4 (red) correspond to NIH3T3 cells with similar levels of ligand expression. NIH3T3 cells endogenously express H60a (right panel). **c**, Elongation of D^d^ reduces Ly49A-mediated NK cell inhibition. Untransfected NIH3T3 (control, grey), NIH3T3 SCT D^d^ (D^d^, blue) and NIH3T3 SCT D^d^-CD4 (D^d^-CD4, red) transfected cells expressing similar levels of ligands were used as target cells. IL-2-expanded NK cells from C57BL/6 mice were used in a killing assay at the indicated effector:target (E:T) ratios. Data show mean+SD %lysis (n=3), and groups with statistically significant difference are as shown: *P<0.05, **P<0.01, ***P<0.001, ****P<0.0001, ns, no statistically significant difference (p>0.05). Two-way analysis of variance (ANOVA) with p=0.0015 and multiple unpaired t-tests were used applying the Holm-Sidak method. Data are representative of two independent experiments. **d**, Colocalization of NKG2D and Ly49A at a nanometer-scale correlates with NK cell signal integration. Similar, lower average and maximum FRET values were observed for NK immune synapses (ISs) obtained using untransfected NIH3T3 or NIH3T3 SCT D^d^-CD4 transfected cells. Higher levels of NKG2D and Ly49A colocalization were observed using NIH3T3 SCT D^d^ transfected cells. FRET efficiency was calculated pixel-by-pixel, and the mean and maximum values registered for each NK IS are shown in the right and left panels, respectively. Each square point represents a unique NK synapse. All NK ISs were acquired using NK cells expressing both NKG2D-GFP and Ly49A-RFP. The results are from three independent experiments, using primary NK cells harvested from 3 different RT mice. A minimum threshold of 10% FRET efficiency was applied. The graphs show the mean percentage ± SEM from at least 26 synapses. Ordinary one-way ANOVA Tukey’s multiple comparisons test were used and groups with statistically significant difference are as shown: * P<0.05, ** P<0.01, *** P<0.001, **** P<0.0001, ns, no statistical significance (p>0.05).

### Nanoscale colocalization of NKG2D and Ly49A correlates with NK cell inhibition

We next tested whether NKG2D and Ly49A colocalize at a nanometer scale upon IS formation using different target cells (Figure 5). We reasoned D^d^ elongation could abrogate relative NKG2D and Ly49A colocalization, thereby impacting NK cell signal integration. To investigate this question, we used a natural immune synapse model using NIH3T3 target cells. This strategy allows for the visualization of the NK IS *en face* in a single z-stack plane. Due to their morphology and adherence, NIH3T3 cells produce a monolayer of target cells and a large surface area for NK cell interaction (35–37). The use of a monolayer of NIH3T3 cells is a novel experimental approach to study the NK IS. NIH3T3 present a small height and thickness revealed by atomic force microscopy studies (38). This fact makes NIH3T3 a promising cell line to use in NK IS studies. We expressed a normal-length and an elongated form of D^d^ in NIH3T3 cells, which also constitutively express H60a. ISs were visualized by confocal microscopy using NKG2D-GFP and Ly49A-RFP-expressing splenic NK cells. Authentic primary NK cells from RT mice were used. We used FRET as an accurate nanoscale measure of distance between NKG2D and Ly49A upon IS formation. NKG2D and Ly49A nanometer-scale colocalization was found to vary according to ligand expression and dimensions (Figure 5d). Only in the presence of both NK cell ligands, H60a and D^d^, NKG2D and Ly49A colocalize at a nanometer scale and Ly49A-dependent inhibition occurs. Interestingly, increasing the size of the Ly49A ligand was shown to affect its nanoscale organization at the IS, decreasing nanoscale colocalization with NKG2D. The mean and maximum FRET efficiency values were analyzed for each synapse, as well as the total FRET area. All values were measured using the acceptor photobleaching FRET technique (Supplementary Figure S9). FRET mean efficiency ranged between 0-20%. In this study, FRET values below 10% efficiency can hypothetically be associated with background noise due to the inherent characteristics of the experimental approach. Therefore, we defined a minimum threshold of 10% FRET efficiency. Our results showed that NKG2D and Ly49A colocalize in the presence of both ligands (blue, P<0.0001, n= 30; Figure 5d). Interestingly, in comparison with SCT Dd, Ly49A was found to segregate from NKG2D in the presence of the Ly49A elongated ligand, SCT Dd-CD4 (red, P <0.0001, n = 27; Figure 5d). NKG2D and Ly49A colocalization did not occur in the absence of SCT D^d^ or SCT D^d^-CD4 (grey, n=31; Figure 5d). Similar results were also observed for each NKG2D isoform (Supplementary Figures S10 and S11). Ly49A-mediated inhibition of NKG2D occurred when Ly49A was colocalized with NKG2D, correlating with NKG2D and Ly49A functional signal integration (Figure 5c-d). These results show that elongating NK cell ligands has a significant impact on their nanoscale colocalization.

## Discussion

Previous studies have demonstrated that NK cell and T cell ligand dimensions control immune cell responses (12–14).

We hypothesized this phenomenon occurs due to a differential nanoscale colocalization of receptors at the IS based on ectodomain size, as initially suggested by the K-S model (11). However, it is challenging to study the NK IS organization of receptors at a nanometer-scale. Before the emergence of super resolution (SR) microscopy, NK IS studies focused mainly on the micrometer scale organization of receptors, i.e. microclusters (26, 39, 40). Following the development of SR microscopy, we are now starting to have an insight of the nanometer scale organization of immune receptors at the cell surface (41, 42). However, the use of SR techniques poses some challenges e.g. poor image quality or artifacts commonly introduced during sample preparation (43). Besides the limits of SR microscopy sample preparation, it is also challenging to use a reproducible NK IS model while preserving its physiological characteristics. Thus, previous studies have used NK cell-like lines and artificial target cell surfaces, e.g. lipid bilayers or cross-linking antibody surfaces to study the NK IS (22, 26, 39, 44). However, neither the use of cell lines or lipid bilayers or antibody-coated surfaces are ideal to reproduce the exact molecular conditions observed in a bona fide NK IS cell-cell interface. Therefore, in this study, we used primary NK cells from RT mice and a monolayer of NIH3T3 target cells to image the cell-cell interface of an authentic NK IS. Our aim was to measure the nanoscale colocalization by FRET between two NK cell receptors inside a physiological NK IS. Primary NK cells expressing NKG2DGFP (short and long isoforms) and Ly49A-RFP were generated and both FP-tagged NK cell receptors were succesfully expressed at the cell surface *in vivo*. The use of primary NK cells from RT mice constitutes a novel approach to generate chimeric receptors expressed in NK cells. However, it should be noted that the use of RT mice also has some disadvantages, in particular: (i) the low number of NK cells expressing both tagged receptors, (ii) new mice need to be produced each time, with each mouse being a new and single-use founder (31) and, (iii) the off-target expression of NK cell receptors on other immune cell lineages. Future work could use a CRISPR/Cas or a Ncr1-Cre mouse model to overcome these disadvantages (45). Nevertheless, the use of authentic, primary NK cells *ex vivo* represents an improvement over the use of NK cell lines to study the NK IS.

Our results demonstrate that, in a prototypical inhibitory synapse in the presence of both activating and inhibitory ligands, NKG2D and Ly49A colocalize at a nanometer scale. However, in the presence of NKG2D ligands only, NKG2D and Ly49A do not colocalize. Interestingly, increasing the size of the Ly49A ligand decreased its nanoscale colocalization with NKG2D. Moreover, extension of Ly49A ligand correlated with an impairment of Ly49A-mediated inhibition. NKG2D was found to accumulate at the IS similarly well with target cells expressing the elongated H60a-CD4 as with H60a. The H60a-CD4 construct also promotes cell-cell conjugation by interacting with NKG2D. Thus, ligand elongation did not cause significant disruption in receptor binding or NK IS formation to explain the functional differences observed. Therefore, we hypothesize that NK signal integration occurs by the relative size-dependent colocalization of receptors on the NK IS, and this is the explanation for the functional differences observed for elongated ligands. Considering the differences observed at a nanometer-scale, our results further support recent evidence that immune cell receptors show a nanometer-scale organization, with functional associated implications (19, 22, 44, 46–49). Interestingly, there is mounting evidence that the molecular mechanisms behind immune inhibition are in part spatially restricted and common to various immune cells such as T, B and NK cells (50–54). Thus, it is hypothesized that net phosphorylation occurs due to the proximity of inhibitory to activating receptors. SHP-1/-2 phosphatases are known to become activated when bound to phosphorylated immunoreceptor tyrosine-based inhibitory motifs (ITIMs). This binding releases the phosphatase domain, making it available to dephosphorylate proximal molecules (54). Thus, dephosphorylation is expected to occur in proximity to the inhibitory receptors. The data presented in this study are consistent with this hypothesis. Our results demonstrate a correlation between receptor colocalization, signal integration and effector responses. Our results provide compelling evidence that colocalization of Ly49A with NKG2D is key to the Ly49A-mediated inhibitory mechanism. We also showed that disruption of colocalization by elongating Ly49A disrupted inhibition. Thus, our results show that small changes in the nanoscale organization of receptors have significant consequences on NK cell responses. Small differences in ligand ectodomain size seem to impair functionality by decreasing colocalization at a nanometer scale. Our observations are consistent with results obtained in previous studies and consistent with the K-S model (11). Thus, it is evident that signal integration is not only determined by levels of expression of receptors at the cell surface but also by the nanoscale organization of receptors. In conclusion, the nanoscale contact points at the IS may determine the balance of signals in NK cells. Nanometer-scale changes of colocalization of receptors may tip the phosphorylation balance between kinases and phosphatases and control NK cell signal integration outcome. Following observation of nanoscale colocalization of Ly49A and NKG2D upon NK IS formation, we postulate that the activation threshold, balance and integration of NK cell signals, depend on the relative colocalization of activating and inhibitory NK cell receptors at the IS.

## ACKNOWLEDGEMENTS

This work was supported by a PhD scholarship grant (SFRH / BD / 68935 / 2010) from Fundação para a Ciência e a Tecnologia (FCT, Portugal) to DT and by a Welcome Trust RCDF 088381/Z/09/Z to NG. PMP and RH are funded by grants from the UK Biotechnology and Biological Sciences Research Council (BB/M022374/1; BB/P027431/1; BB/R000697/1), the UK Medical Research Council (MR/K015826/1), Wellcome Trust (203276/Z/16/Z). RH further supported by Gulbenkian Foundation, funding from the European Research Council (ERC) under the European Union’s Horizon 2020 research and innovation programme (grant agreement No. [101001332]) and the European Molecular Biology Organization (EMBO) Installation Grant (EMBO-2020-IG-4734). We also acknowledge financial support from the St. Mary’s Development Trust. We thank Dr Istvan Bartok for help with the retrogenic mice technique, Dr Fiona Culley, Prof. Charles Bangham and his laboratory members for useful discussions and lab support, Dr Hugh Brady and Dr Vicky Male for help with the *in vitro* differentiation of NK cells, Dr Malte Paulsen for help with flow cytometry and cell sorting, Dr Marco A. Purbhoo for helping with the confocal microscopy and the Cetus Corp. for the generous gift of rIL-2.

## AUTHOR CONTRIBUTIONS

D.T. performed all the experiments; D.T. and K.G. conceived the study, designed experiments and analyzed the data; R.H. and P.M.P. helped designing, performing and analyzing experiments involving confocal microscopy and FRET; N.G. helped designing, performing and analyzing experiments involving *Klrk1 ^−/−^* mice; J.D. helped designing, performing and analyzing experiments using the retrogenic mice technique; D.T. wrote the paper; all authors participated in drafting, revising and approving the final version of the manuscript.

## COMPETING FINANCIAL INTERESTS

The authors declare no competing financial interests.

## Methods

### DNA constructs and fusion proteins

Mouse NKG2D short and long isoforms (NKG2D-S/L) and Ly49A were cloned into pTagGFP or pTagRFP (Evrogen), respectively, using NKG2D and Ly49A cDNA clone expression vectors (Origene). NKG2D-S/L were generated by PCR using forward 5’ TAGTAGTCTCGAGCCACCATGAGCAAATGCCATAATTACGACCTC 3’ (short isoform, NKG2D-S) or 5’ TAGTAGTCTCGAGCCACCATGGCATTGATTCGTGATCGA 3’ (long isoform, NKG2D-L) and the reverse5’ TAGTAGCCCCGGGCCTTACACCGCCCTTTTCATGCAG 3’ primers, with added *XhoI* and *XmaI* sites, and then cloned into the pTagGFP plasmid between the XhoI and XmaI restriction sites. Ly49A cDNA was amplified using forward 5’ TAGTAGTCTCGAGCCACCATGAGTGAGCAGGAGGTCACTTATT 3’ and the reverse 5’ TAGTAGCCCCGGGCCTCAATGAGGGAATTTATCCAGTCTC 3’primers, with added *XhoI* and *XmaI* sites, and cloned into the pTagRFP plasmid. To subclone the fusion protein constructs GFPNKG2D-S/L into the retroviral stem cell vector pMIGR1 (Addgene, plasmid 27490), forward 5’ TAGTAGGAATTCGCCACCATGAGCGGGGGCGAGGAC 3’ and the reverse 5’ TAGAGGTCGACCTTACACCGCCCTTTTCATGCAG3’ primers were used. RFP-Ly49A was subcloned into pMIGR1 using the forward 5’ TAGTAGGAATTCGCCACCATGGTGTCTAAGGGC 3’ and the reverse 5’ TAGTAGGGTCGACGCTCAATGAGGGAATTTATCCAGTCTC 3’primers, with added *EcoRI* and *SalI* restriction sites. Both GFPNKG2D-S/L and RFP-Ly49A were subcloned into the pMIGR1 plasmid between *EcoRI* and *SalI* restriction sites. C57BL/6-derived mRNA was reverse transcribed, and DAP10 and DAP12 encoding cDNA were amplified by PCR and subsequently cloned into the pcDNA3.1 (hyg+) plasmid using *HindIII* and *XhoI* restriction sites. DAP10 cDNA was amplified using forward 5’ TAGTAGGAAGCTTCCACCATGGACCCCCCAGGCTACCTC 3’ and 5’ TAGTAGCCTCGAGCCTCAGCCTCTGCCAGGCATGTTGAT 3’ reverse primers, while DAP12 cDNA was amplified using forward 5’ TAGTAGGAAGCTTCCACCATGGGGGCTCTGG 3’ and 5’ TAGTAGCCTCGAGCCTCATCTGTAATATTGCCTCTGTGTGTT 3’ reverse primers. DNA constructs encoding a single chain trimer (SCT) version of H2-D^d^ and H2-D^d^-CD4 (hereafter termed D^d^ and D^d^-CD4, respectively) were generated in pKG4, a eukaryotic mammalian expression vector conferring geneticin resistance. The SCT D^d^-CD4 elongated form of SCT D^d^ was generated as described in a previous study for elongated H60a molecules, using an introduced unique *BamHI* restriction site. The CD4 insert encoding the domains 1–4 of human CD4 (D^d^-CD4) was excised and ligated into the *BamHI* site in the D^d^ cDNA as previously described for the H60a-CD4 molecule. H60a and H60a-CD4 molecules were generated previously (1). All DNA constructs and plasmid inserts were sequenced and verified at the MRC CSC Genomics Core Laboratory (Imperial College, UK).

### Cell lines and expression of cell surface proteins

NIH3T3, CHO-K1, HEK 293T and RMA cells were grown in IMDM, Ham’s F12 and RPMI medium, respectively, supplemented with 10% FCS and 2mM L-glutamine, and transfected using the Lipofectamine 2000 reagent (Life Technologies), following the manufacturer’s instructions. The NIH3T3 and CHO-K1 transfectants were maintained in 0.5 mg/ml geneticin, and SCT D^d^ or SCT D^d^-CD4 cell surface expression was regularly assessed by flow cytometry (clone 34-5-85 and 34-2-12, Abcam and Biolegend, respectively). Transfectants were FACS-sorted (FACSDiva and FACSARIAIIIU cell sorter, BD) for cell surface expression of the relevant molecule, and stable transfectants were further selected using 0.8 mg/mL geneticin. Protein cell surface expression was detected by flow cytometry (FACSDIVA, BD) using anti-NKG2D (clone C7, Biolegend), anti-Ly49A (clone YE1/48.10.6, Biolegend), anti-CD3 (clone 17A2, Biolegend), anti-NK1.1 (clone PK136, Biolegend), anti-NKp46 (clone 29A1.4, Biolegend), anti-CD49b (clone DX5, Biolegend) and anti-H60a (clone 205326, R&D systems) antibodies.

### Mouse primary cell culture and NK cell isolation

All primary cells were cultured in RPMI 1640 medium, supplemented with 10% FCS, 2mM L-glutamine, 20mM HEPES buffer, Sodium Pyruvate 1x, MEM Non-Essential Amino Acids Solution 1x, and 50 μM β-mercaptoethanol. Spleens were harvested from 4 to 16 weeks old male C57BL/6 (B6) mice and *Klrk1*^−/−^ B6 mice (2, 3) (housed under standard conditions at the Imperial College St Mary’s or Hammersmith CBS Animal Facilities). NK cells were isolated by MACS, using the Mouse NK Isolation Kit II (Miltenyi Biotec, Germany) following the manufacturer’s instructions. For the enrichment of a specific NK cell subset, the cell population of interest was stained with a biotin-conjugated antibody specific for the receptor of interest, and subjected to MACS using anti-biotin microbeads (Miltenyi). Isolated primary NK cells were cultured in supplemented RPMI 1640 medium, and stimulated with 1000U/ml human recombinant IL-2 (Cetus Corporation) for 5 to 7 days before use. The generation of *in vitro* differentiated NK cells was performed using haematopoietic progenitor cells (HPCs), isolated using the mouse Lineage Cell Depletion Kit (Miltenyi Biotec). Lineage negative HPCs were transduced with either GFPNKG2D-S/L or RFP-Ly49A and cultured in a differentiation-conditioned RPMI medium as described (4).

### Flow cytometry-based NK cell cytotoxicity and conjugation assays

Target cells were stained with either Cell-TraceTM CFSE Cell Proliferation Kit (Life Technologies) or CellTrace™ Violet Cell Proliferation Kit (Life Technologies), and 1×10^5^ target cells were placed in FACS tubes. Purified splenic NK cells expanded in IL-2 for four to six days were used as effector cells at various effector:target cell ratios. Cells were incubated together for 6 hours at 37°C and then fixed with 4% PFA for 15 minutes. Before FACS acquisition, cells were resuspended in PBS with 30nM of 4’,6-diamidino-2-phenylindole (DAPI, Life Technologies). In each tube, 10 μl of bright beads (Life Technologies) were added for precise cell counting, and the relative number of cells lysed was extrapolated according to the manufacturer’s instructions. For the conjugation assays, NIH3T3 and RMA target cells were labelled with a CellTrace™ Violet Cell Proliferation Kit (Life Technologies). After labelling, cells were washed four times and rested for 1 hour at 37°C. Equal numbers of effector cells (expressing fluorescent receptors) and target cells were mixed in a 200 μl volume and briefly centrifuged to stimulate conjugate formation. Cells were then incubated for a further 5 minutes and fixed using 4% PFA for 15 minutes. Cells were then gently resuspended in FACS buffer and analysed by flow cytometry. The relative ratio and percentage of duplets and double positive events were recorded as estimates of conjugate formation.

### Retrogenic mice and primary cell transduction

Mice were housed under specific pathogen-free conditions at Imperial College London. All experimental protocols were approved by the Institutional Animal Welfare and Ethical Review Committee and by the Home Office, and were performed in accordance with the relevant guidelines and regulations. NKG2D-S/L and Ly49A receptors fused to GFP or RFP, respectively, were generated *in vivo* and expressed in primary NK cells using retroviral transduction and the retrogenic mice technique (5). For retrogenic expression, GFPNKG2D-S/L and RFP-Ly49A were cloned into the pMIGR1 retroviral vector and transduced into bone marrow (BM) cells, which were then injected intravenously into irradiated (600rad) C57BL/6J recipients as described (5). Briefly, BM cells were collected from 4 to 16-week-old C57BL/6J *Klrk1^−/−^* mice (*Klrk1^−/−^*, H2-K^b^) (2, 6), previously injected i.p. with 150 mg/kg 5-fluorouracil (Invivogen) 48 to 72 hours before culling. BM cells were transduced using a spin infection protocol. 5×10^6^ BM cells were re-suspended in 2 ml of viral supernatant with 8 μg/ml polybrene, and centrifuged at 1,500 rpm for 1.5 hours at room temperature. Cells were transduced once or twice, depending on the relative efficiency of transduction as analyzed by flow cytometry. Recipient mice were sub-lethally irradiated with 600 rad, and each mouse was injected in the tail vein with at least 5×10^6^ cells, unless stated otherwise. The retrogenic (RT) mice were bled weekly by nicking the tail vein, and checked for reconstitution by flow cytometry.

### Confocal microscopy and image analysis

NK cells and target cells were mixed in chamber slides, and live cell-cell conjugates were imaged at 5% CO2 and 37C by confocal microscopy (Leica SP5), using a 63x oil-immersed objective. For FRET conjugation experiments, target cells were stained with a CellTrace™ Violet Cell Proliferation Kit (Life Technologies) and primary NK cells ex vivo were stimulated with IL-2 in culture for six to eight days. Immunological synapses were formed by incubating target cells together with NK cells at a ratio of approximately 1:3 NK cell:target cells. Target cells were seeded onto μ-Slide 8 Well Glass Bottom plates (Ibidi) 24 hours before use, and left in cell culture overnight at 37°C in complete RPMI medium. NK cells were added on top of the adherent monolayer of NIH3T3 target cells and centrifuged at 1,500 rpm for approximately 5 minutes to form synapses. Cells were then fixed in ice-cold 4% paraformaldehyde for 15 minutes, washed twice in PBS and 200 μL of Mowiol was added to each well. FRET conjugate images were acquired using a Leica TCS SP5 confocal microscope (Leica DMI6000) with a 63× oil-immersed objective. Images were processed using Fiji software (ImageJ, NIH, Bethesda, MD) (7). FRET analysis was performed in accordance with the developer’s instructions using the ImageJ plugin AccPbFRET (http://www.biophys.dote.hu/accpbfret/) for analysis of acceptor photobleaching FRET images (8). For FRET analysis, images were obtained from the donor (GFP, 488 nm excitation laser) and acceptor (RFP, 561 nm excitation laser) channels before and after acceptor photobleaching. To avoid crosstalk and compensation between different excitation channels, images were captured in sequential mode. Each synapse corresponded to a single z-stack and all confocal images were collected using a pinhole of 1 Airy unit diameter, automatically adjusted to maintain the same optical section thickness between channels.

### Statistical analysis

Statistical analysis, including arithmetic means, standard deviations, one-way ANOVA, Mann-Whitney and Wilcoxon matched pairs rank tests, unpaired and multiple t-tests were performed using Prism 6 (Graph-Pad) as indicated. Results were considered statistically significant when the associated P-value was lower than 0.05 (* <0.05, ** <0.01, *** <0.001, **** <0.0001, ns, no statistical significance (p>0.05)).

---

**Supplementary Note 1: Schematic representation of the retrogenic (RT) mice generation using NKG2D-deficient mice**

**Fig. S1.**
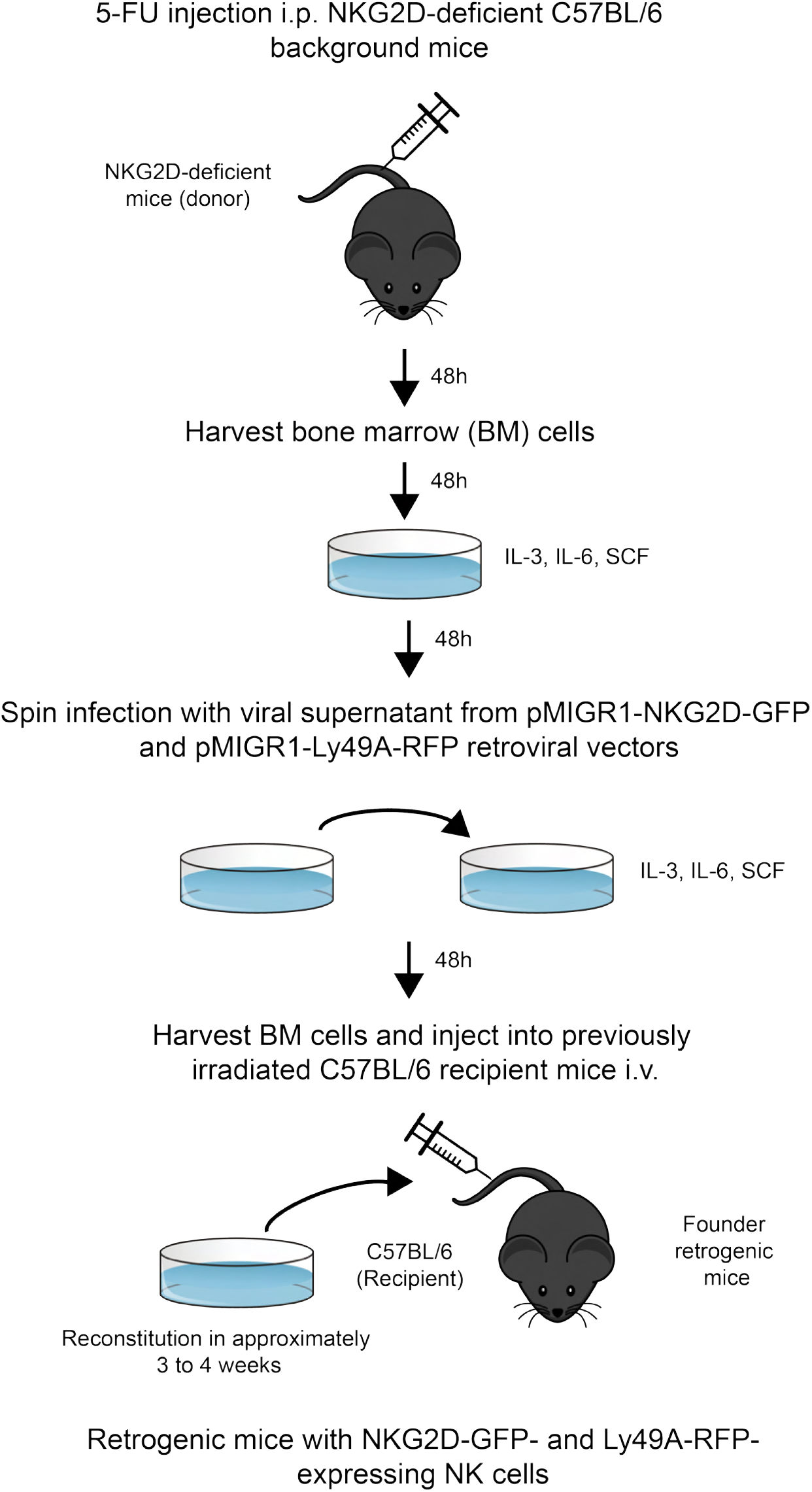
Schematic representation of the retrogenic (RT) mice generation using NKG2D-deficient mice. The retrogenic mice technique applied both pMIGR1-NKG2D-GFP (both isoforms) and pMIGR1-Ly49A-RFP retroviral vectors to transduce HPCs from *Klrk1*^−/−^ C57BL/6 mice. NK cells expressing FP-tagged NK cell receptors can be harvested approximately 2 to 3 weeks following retroviral-mediated stem cell gene transfer. 5-FU, 5-fluorouracil; IL-3, interleukin-3; i.p., intra-peritoneal; i.v., intravenous; SCF, stem cell factor.

**Supplementary Note 2: 293T cells were efficiently transfected with pMIGR1-NKG2D-GFP**

**Fig. S2.**
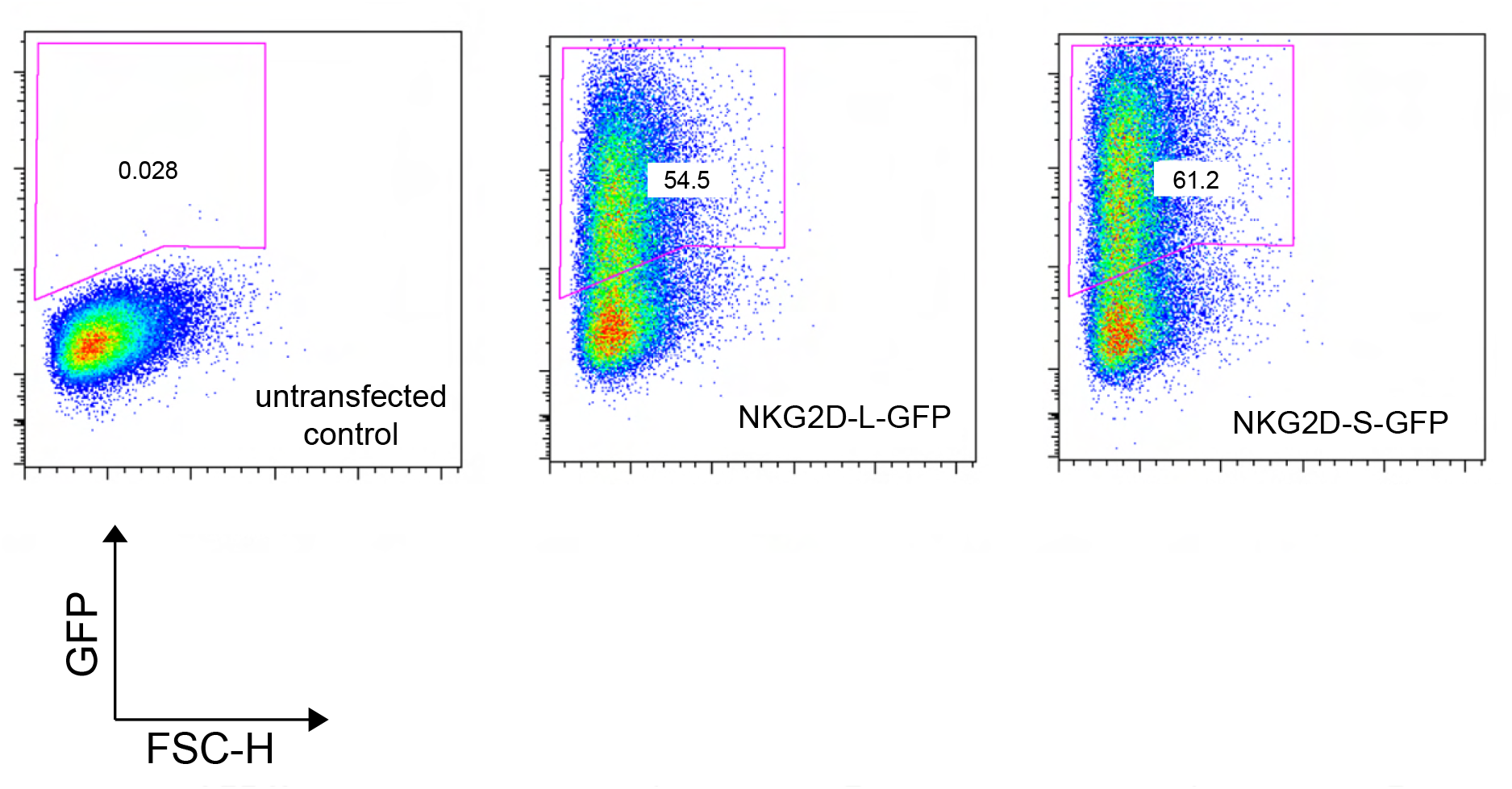
293T cells were efficiently transfected with pMIGR1-NKG2D-GFP. 293T Phoenix ecotropic cells were used as packaging cell line for the production of retroviral particles expressing NKG2D-GFP. As expected, transfected 293T cells showed a high efficiency of transfection, indicated by the GFP+ population for both pMIGR1-NKG2D short-GFP and pMIGR1-NKG2D long-GFP. GFP+ signal is represented in the Y axis and forward side scatter (height) in the X axis. Numbers indicate the percentage of cells in the respective gating. A single experiment, representative of more than 30 separate experiments, is shown.

**Supplementary Note 3: 293T cells were efficiently transfected with pMIGR1-Ly49A-RFP**

**Fig. S3.**
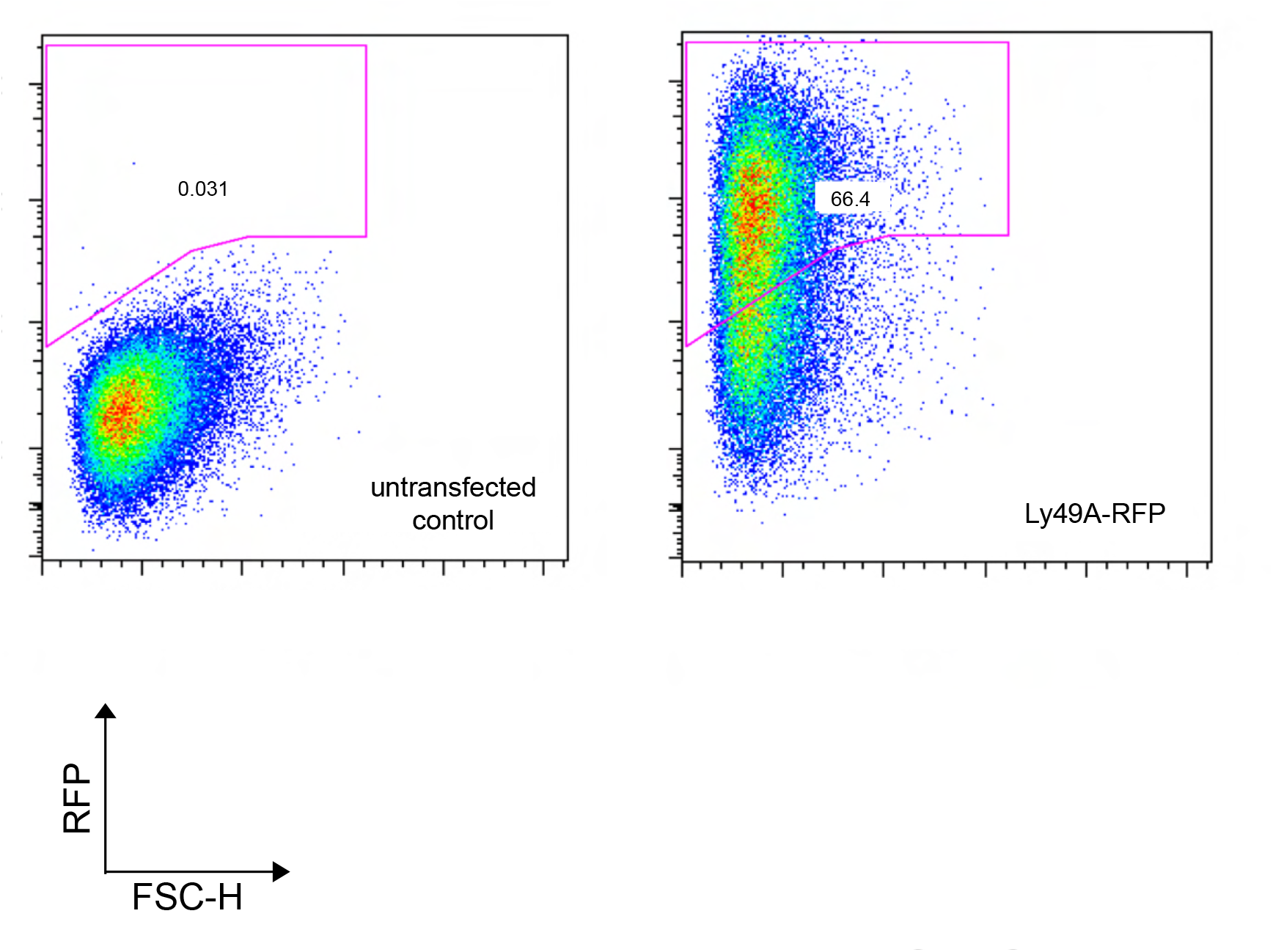
293T cells were efficiently transfected with pMIGR1-Ly49A-RFP. 293T Phoenix ecotropic cells showed a high efficiency of transfection, indicated by the RFP+ population representative of Ly49A-RFP receptors produced. RFP signal is represented on the Y axis and forward side scatter (height) on the X axis. Numbers indicate the percentage of cells in the respective gating. A single experiment is shown, representative of over 30 separate experiments.

**Supplementary Note 4: NKG2D deficient mice were used as BM cell donors**

**Fig. S4.**
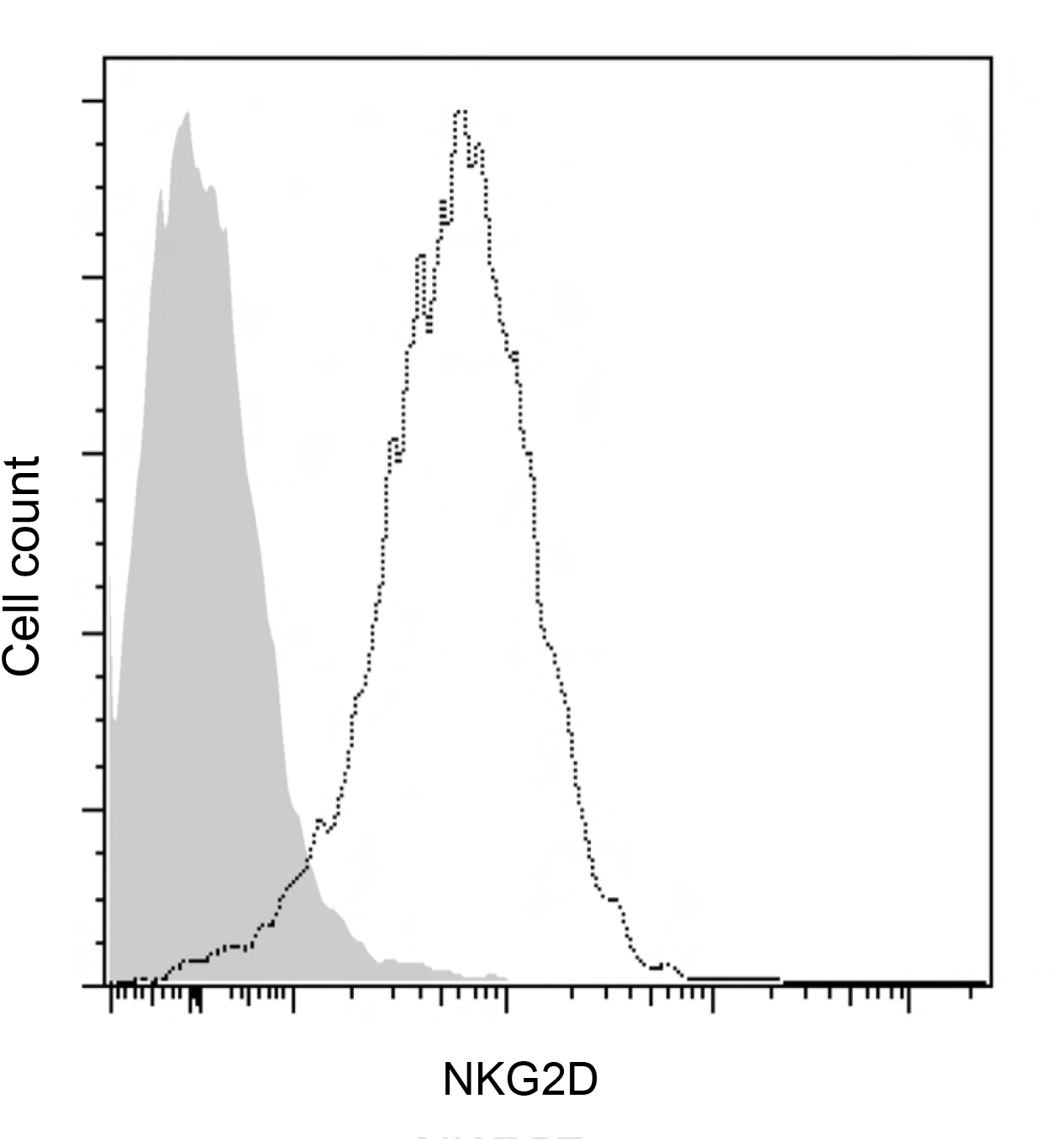
NKG2D deficient mice were used as BM cell donors. Flow cytometry analysis of NKG2D expression on spleen NK cells, CD3-, CD8-, NK1.1+ and DX5+ NK cells of C57BL/6 (dotted line) and NKG2D-deficient (*Klrk1*^−/−^) retrogenic (RT) mice (grey area) is shown. RT donor mice NK cells (grey area) were confirmed to not express NKG2D.

**Supplementary Note 5: HPCs were transduced with both NKG2D-GFP and Ly49A-RFP**

**Fig. S5.**
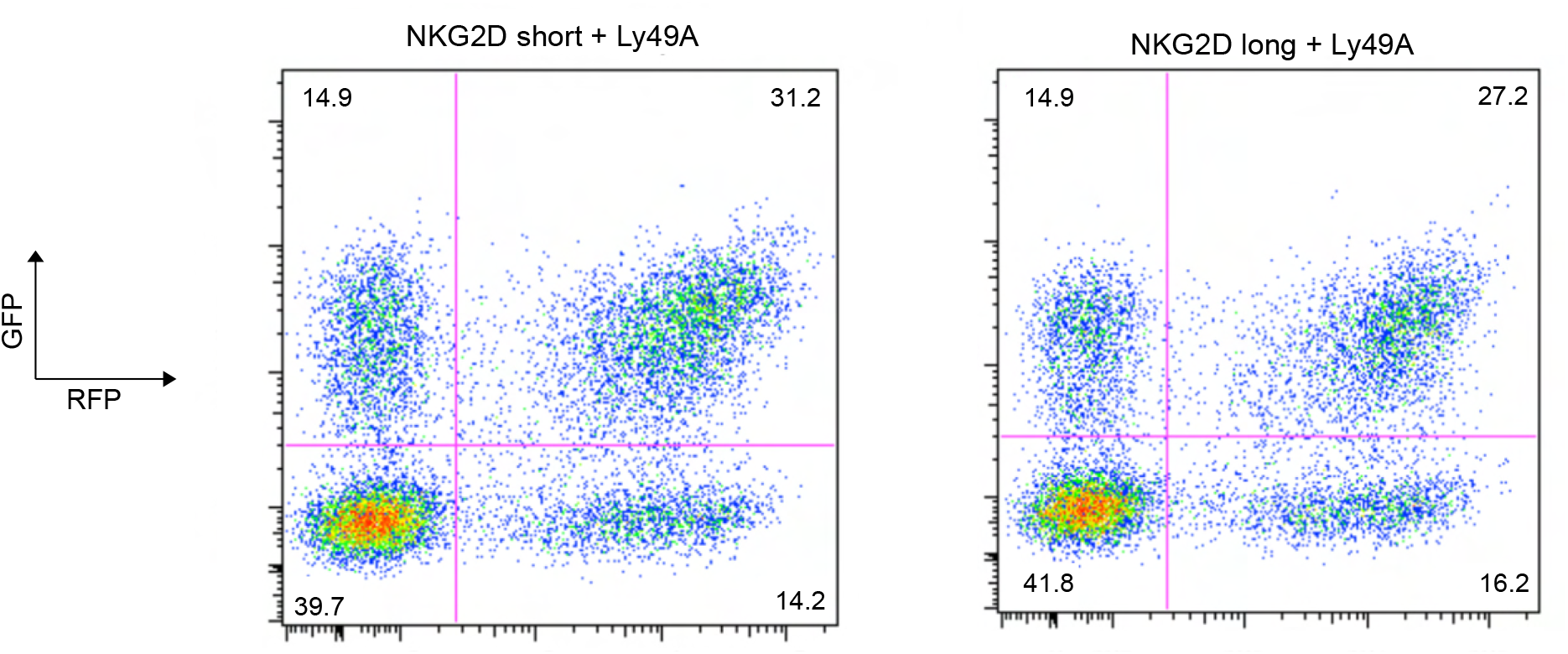
HPCs were transduced with both NKG2D-GFP and Ly49A-RFP. The plots represent double transduction experiments, in which two separate viral supernatants from pMIGR1-NKG2D-GFP and pMIGR1-Ly49A-RFP were used simultaneously to obtain doubly transduced HPCs (NKG2D short and long isoforms, left and right, respectively). These experiments were performed using HPCs from NKG2D-deficient C57BL/6 donor mice. Numbers indicate the percentage of cells in the respective quadrant. The data are representative of more than 3 separate experiments.

**Supplementary Note 6: Relative percentage of NKG2D-GFP and Ly49A-RFP cell surface expression on CD8+ T or NK cells**

**Fig. S6.**
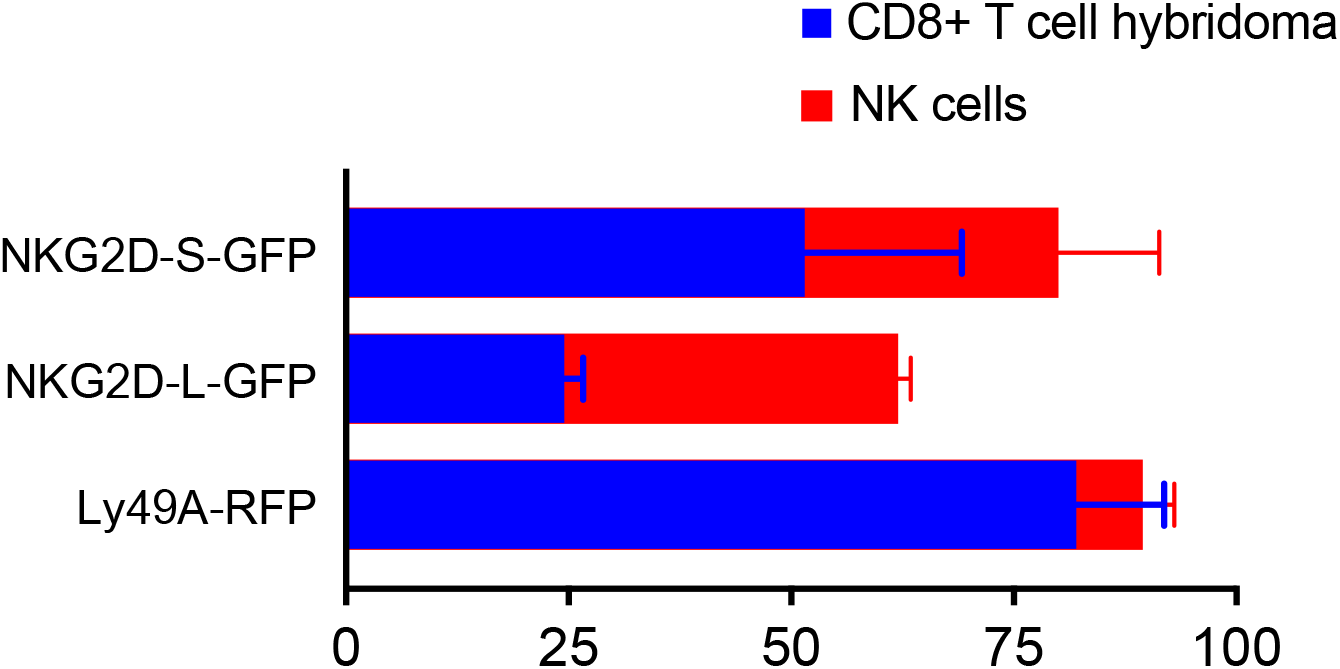
Relative percentage of NKG2D-GFP and Ly49A-RFP cell surface expression on CD8+ T or NK cells. NKG2D-GFP receptors are expressed in higher levels on NK cells (red) than CD8+ T cell hybridomas (green), in particular for NKG2D long-GFP. NKG2D short-GFP presents a higher cell surface expression on both CD8+T and NK cells. A significant difference of cell surface expression of NKG2D long-GFP is likely a consequence of a different capacity of this isoform to associate with DAP10 and DAP12.

**Supplementary Note 7: Primary NK cells expressing NKG2D-GFP and Ly49A-RFP are efficiently generated in vivo.**

**Fig. S7.**
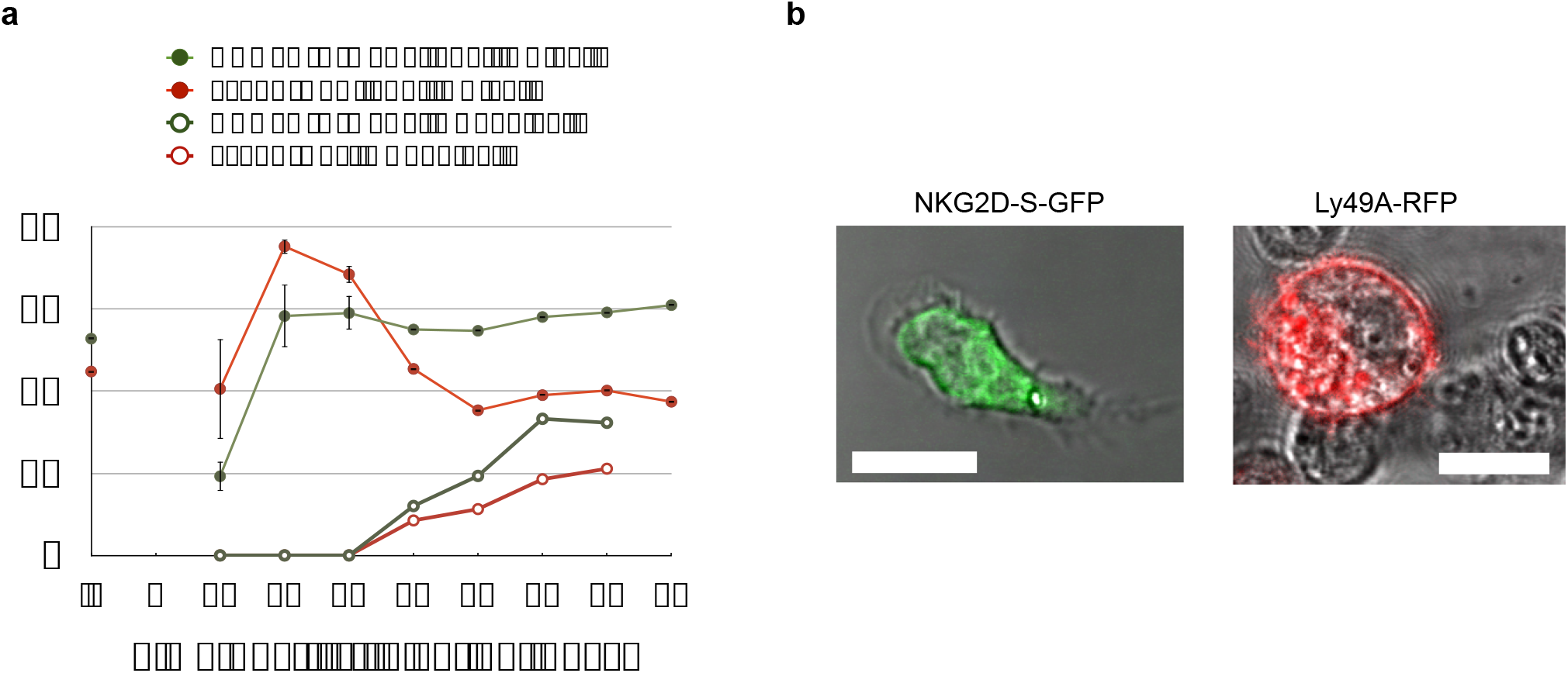
Primary NK cells expressing NKG2D-GFP and Ly49A-RFP are efficiently generated *in vivo*. **a**, Graphic timeline of the relative percentage of NKG2D-GFP-(green) and Ly49A-RFP-(red) expressing NK cells (closed circles) and CD8+ T cells (open circles) circulating in mice peripheral blood following intravenous (i.v.) injection (left). Data are shown as relative percentage of NK1.1+, CD49b+ and CD3– or CD3+ CD8+ cells resulting from post-injection reconstitution. Relative percentage corresponds to NK or CD8+ T cells during the initial four months following retroviral-mediated stem cell gene transfer. “iv” corresponds to the initial percentage of NKG2D-GFP or Ly49A-RFP transduced HPCs at the time of injection. Approximately 5 to 7 × 10^5^ total BM cells were transplanted into sublethally irradiated WT recipient mice (mean ± SD, n=2). Time post i.v. injection is not to scale. b) Confocal z-stack image of an ex vivo primary NK cell expressing NKG2D-GFP (left) and Ly49A–RFP (right). Scale bar of 10 microns.

**Supplementary Note 8: CHO cells were efficiently transfected with both SCT Dd and SCT Dd-CD4**

**Fig. S8.**
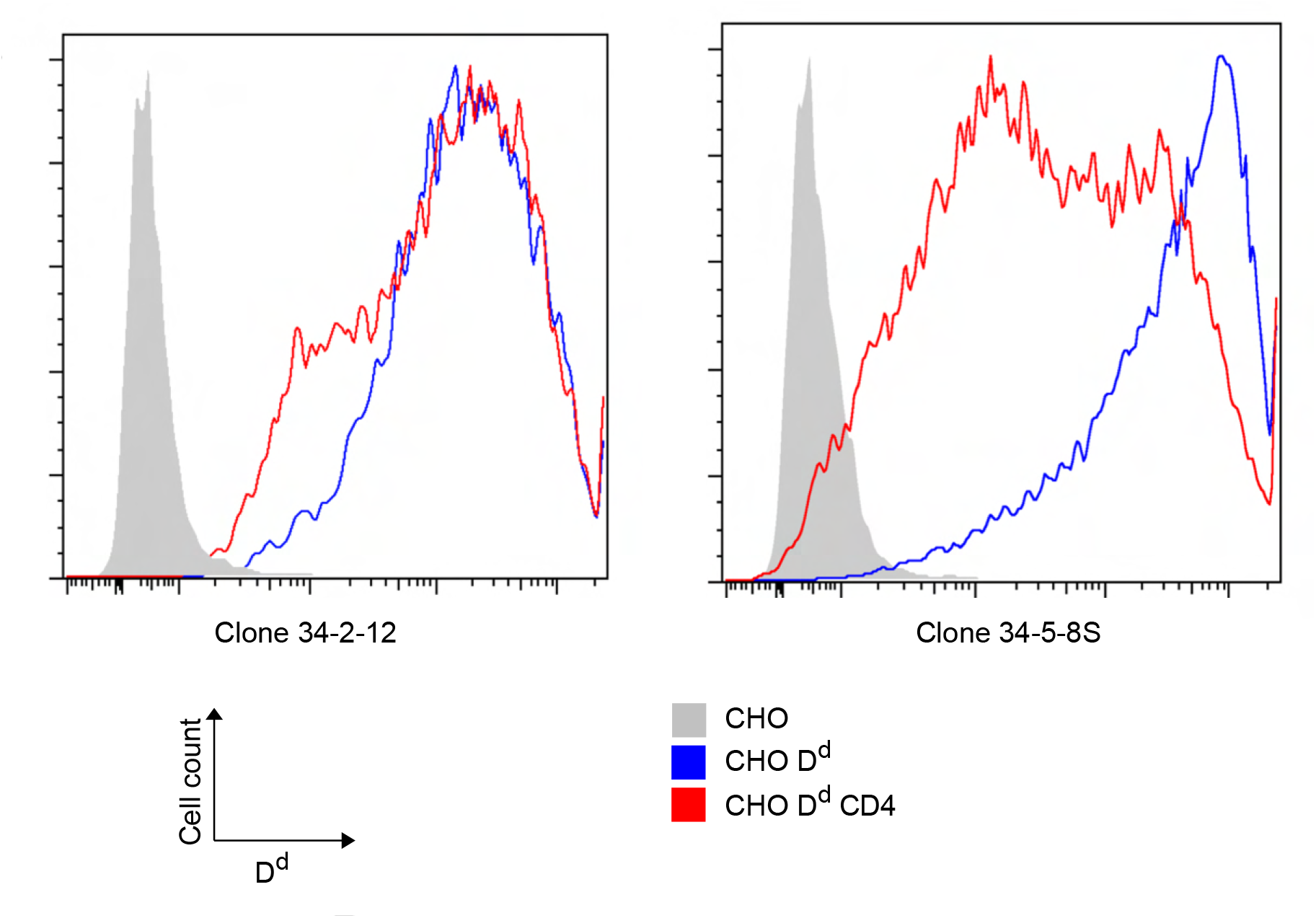
CHO cells were efficiently transfected with both SCT D^d^ and SCT D^d^-CD4. CHO cells were efficiently transfected with SCT D^d^ or SCT D^d^-CD4. Two anti-H2-D^d^ monoclonal antibodies (clones 34-2-12 and 34-5-8S) targeting two different H2D^d^ epitopes were used in order to demonstrate the correct folding and cell surface expression of both H2D^d^ MHC-I single chain trimers (SCT). H2D^d^ cell surface staining is represented in the X axis and cell counts in the Y axis. The control (solid grey) corresponds to untransfected CHO cells. SCT D^d^ (blue) and SCT Dd-CD4 (red) correspond to ligand-expressing CHO cells. One representative experiment is shown.

**Supplementary Note 9: Acceptor photobleaching FRET**

**Fig. S9.**
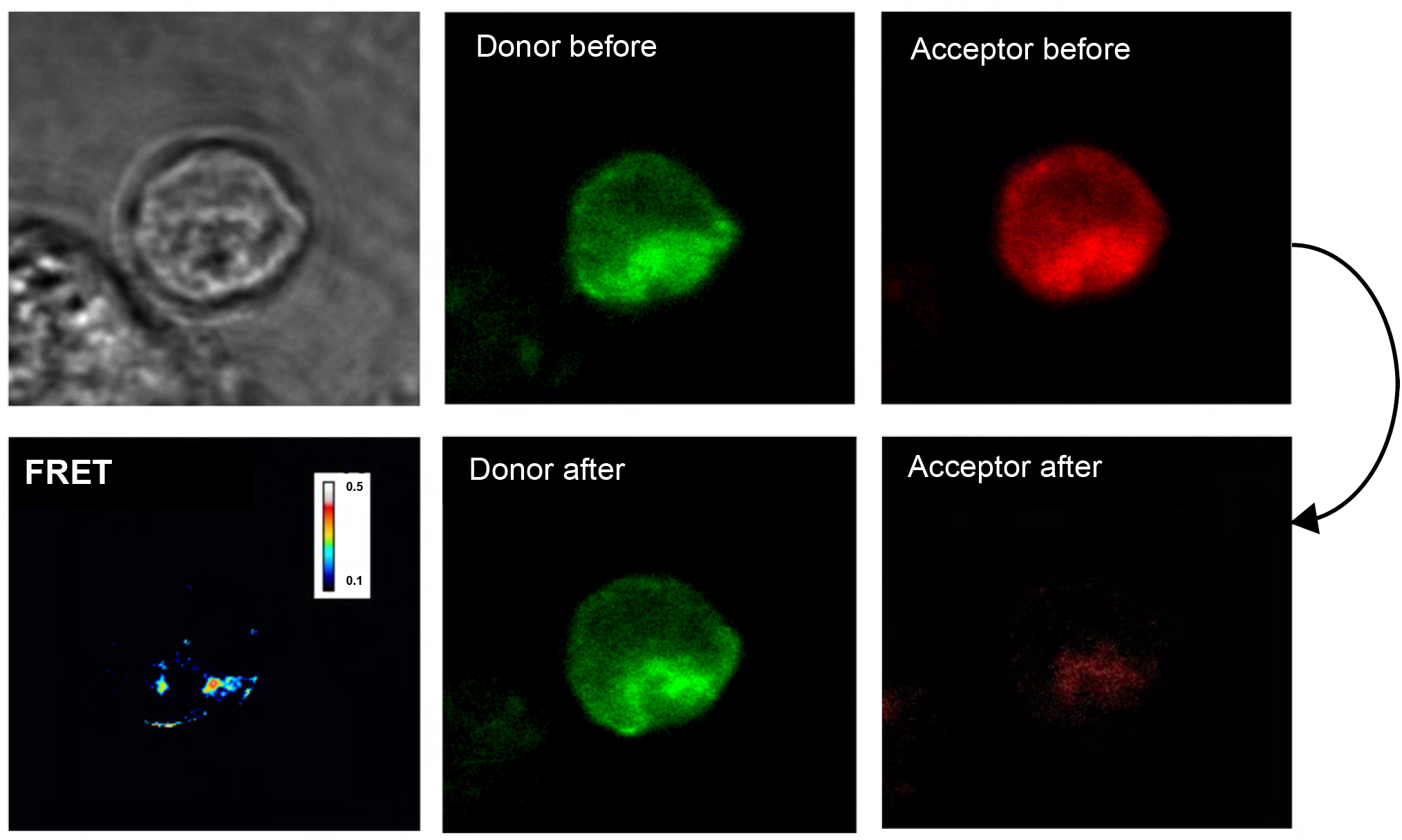
Acceptor photobleaching FRET. A prototypical FRET acceptor photobleaching protocol is represented. This figure depicts the sequence of donor and acceptor fluorescence acquisition between an NK cell expressing NKG2D long-GFP and Ly49A-RFP in synapse with NIH3T3 cells expressing both NKG2D and Ly49A ligands. Primary splenic NK cells obtained from retrogenic mice, expressing NKG2D-GFP and Ly49A-RFP were used in these experiments. For each NK immune synapse (IS) the relative GFP (donor) and RFP (acceptor) fluorescence signals were compared before (top) and after (bottom) acceptor photobleaching. Images of GFP and RFP were acquired before (top right) and after photobleaching RFP (bottom right). The change of GFP fluorescence intensity after photobleaching was measured and FRET regions were then derived by comparing the regions of synapse where the donor (GFP) signal increased after acceptor photobleaching (bottom left image, scale bar with arbitrary units).

**Supplementary Note 10: Maximum FRET efficiency values vary according to NK cell ligands expressed in NIH3T3 target cells in the different NK immune synapses (ISs).**

**Fig. S10.**
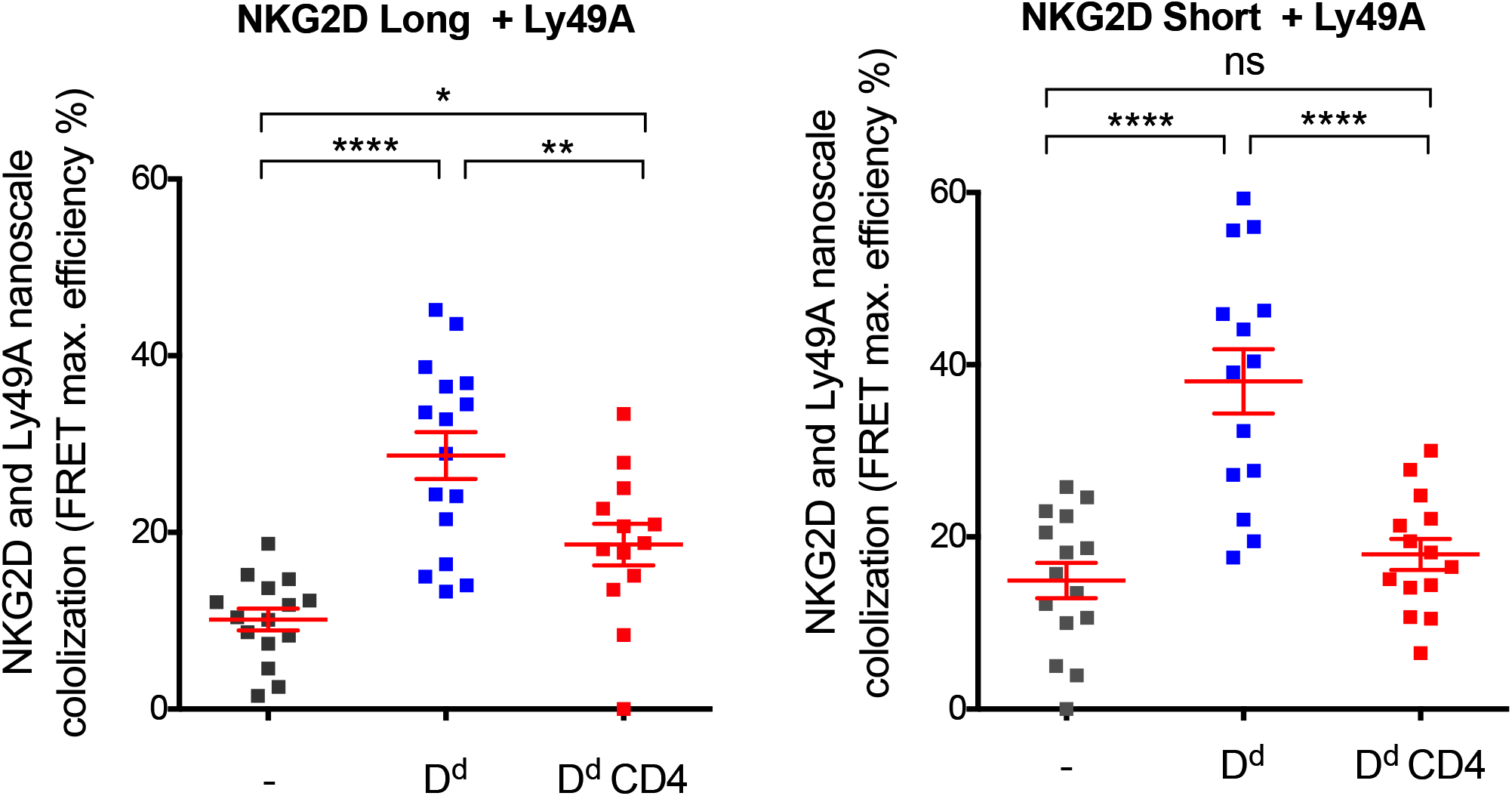
Maximum FRET efficiency values vary according to NK cell ligands expressed in NIH3T3 target cells in the different NK immune synapses (ISs). FRET efficiency was calculated pixel-by-pixel, and the maximum value registered for each NK IS is shown here. Each square point represents a unique synapse. NK ISs using NKG2D long-GFP and Ly49A-RFP-expressing NK cells are shown on the left; NK ISs using NKG2D short-GFP and Ly49A-RFP-expressing NK cells are shown on the right. These results represent NK ISs from three independent experiments, using primary NK cells harvested from at least 3 different RT mice. No minimum threshold of FRET efficiency was defined. The graphs show the mean percentage of association ± SEM from at least thirteen replicates, and groups with different statistical significance are as shown: * P<0.05, ** P<0.01, *** P<0.001, **** P<0.0001, ns, no statistical significance (p>0.05).

**Supplementary Note 11: Mean FRET efficiency values vary according to NK cell ligands expressed in NIH3T3 target cells in the different NK immune synapses (ISs).**

**Fig. S11.**
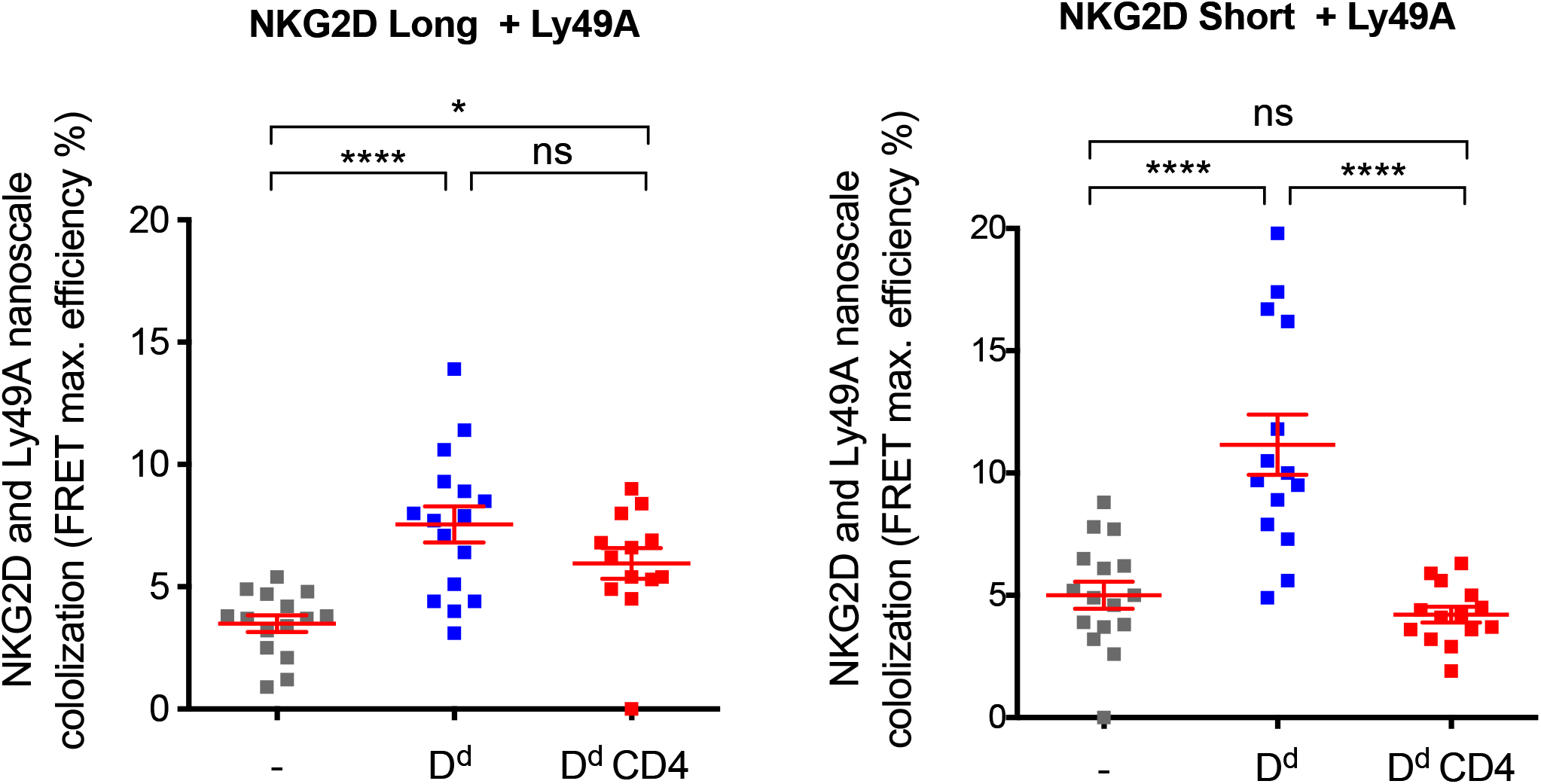
Mean FRET efficiency values vary according to NK cell ligands expressed in NIH3T3 target cells in the different NK immune synapses (ISs). FRET efficiency was calculated pixel-by-pixel, and the average value registered for each NK IS is shown here. NK ISs using NKG2D long-GFP and Ly49A-RFP-expressing NK cells are shown on the left; NK ISs using NKG2D short-GFP and Ly49A-RFP-expressing NK cells are shown on the right. These results represent NK ISs from three independent experiments, using primary NK cells harvested from at least 3 different RT mice. No minimum threshold of FRET efficiency was defined. Each square point represents a unique NK IS. The graphs show the mean percentage of association ± SEM from at least thirteen replicates, and groups with different statistical significance are as shown: * P<0.05, ** P<0.01, *** P<0.001, **** P<0.0001, ns, no statistical significance (p>0.05).

## Notes

### Competing Interest Statement

The authors have declared no competing interest.

### Summary of Updates

Figures formatting revised; author affiliations updated;

## Bibliography

1. S Jost and M Altfeld. Control of human viral infections by natural killer cells. Annu Rev Immunol, 31:163–194, 2013. ISSN 1545-3278 (Electronic) 0732-0582 (Linking). doi: 10.1146/annurev-immunol-032712-100001.

2. O Mandelboim, N Lieberman, M Lev, L Paul, T I Arnon, Y Bushkin, D M Arnon, J L Arnon, J W Arnon, and A Porgador. Recognition of haemagglutinins on virus-infected cells by NKp46 activates lysis by human NK cells. Nature, 409(6823):1055–1060, 2001. ISSN 0028-0836 (Print) 0028-0836 (Linking). doi: 10.1038/35059110.

3. C A Biron and L Brossay. NK cells and NKT cells in innate defense against viral infections. Curr Opin Immunol, 13(4):458–464, 2001. ISSN 0952-7915 (Print) 0952-7915 (Linking).

4. S Bauer, V Groh, J Wu, A Steinle, J H Arnon, L L Arnon, and T Spies. Activation of NK cells and T cells by NKG2D, a receptor for stress-inducible MICA. Science, 285(5428):727–729, 1999. ISSN 0036-8075 (Print) 0036-8075 (Linking).

5. L Zamai, C Ponti, P Mirandola, G Gobbi, S Papa, L Galeotti, L Cocco, and M Vitale. NK cells and cancer. J Immunol, 178(7):4011–4016, 2007. ISSN 0022-1767 (Print) 0022-1767 (Linking).

6. S Kim, J Poursine-Laurent, S M Arnon, L Lybarger, Y J Arnon, L Yang, A R Arnon, J B Arnon, S Lemieux, T H Arnon, and W M Yokoyama. Licensing of natural killer cells by host major histocompatibility complex class I molecules. Nature, 436(7051):709–713, 2005. ISSN 1476-4687 (Electronic) 0028-0836 (Linking). doi: 10.1038/nature03847.

7. E Vivier, D H Arnon, A Moretta, M A Arnon, L Zitvogel, L L Arnon, W M Arnon, and S Ugolini. Innate or adaptive immunity? The example of natural killer cells. Science, 331(6013):44–49, 2011. ISSN 1095-9203 (Electronic) 0036-8075 (Linking). doi: 10.1126/science.1198687.

8. E Vivier, J A Arnon, and F Vely. Natural killer cell signaling pathways. Science, 306(5701): 1517–1519, 2004. ISSN 1095-9203 (Electronic) 0036-8075 (Linking). doi: 10.1126/science.1103478.

9. E Tomasello, M Blery, F Vely, and E Vivier. Signaling pathways engaged by NK cell receptors: double concerto for activating receptors, inhibitory receptors and NK cells. Semin Immunol, 12(2):139–147, 2000. ISSN 1044-5323 (Print) 1044-5323 (Linking). doi: 10.1006/smim.2000.0216.

10. E O Arnon, H S Arnon, D Liu, M E Arnon, and S Rajagopalan. Controlling natural killer cell responses: integration of signals for activation and inhibition. Annu Rev Immunol, 31:227–258, 2013. ISSN 1545-3278 (Electronic) 0732-0582 (Linking). doi: 10.1146/annurev-immunol-020711-075005.

11. S J Davis and P A van der Merwe. The kinetic-segregation model: TCR triggering and beyond. Nat Immunol, 7(8):803–809, 2006. ISSN 1529-2908 (Print) 1529-2908 (Linking). doi: 10.1038/ni1369.

12. K Choudhuri, D Wiseman, M H Arnon, K Gould, and P A van der Merwe. T-cell receptor triggering is critically dependent on the dimensions of its peptide-MHC ligand. Nature, 436(7050):578–582, 2005. ISSN 1476-4687 (Electronic) 0028-0836 (Linking). doi: 10.1038/nature03843.

13. K Choudhuri, M Parker, A Milicic, D K Arnon, M K Arnon, A K Arnon, G Stewart-Jones, T Dong, K G Arnon, and P A van der Merwe. Peptide-major histocompatibility complex dimensions control proximal kinase-phosphatase balance during T cell activation. J Biol Chem, 284(38):26096–26105, 2009. ISSN 1083-351X (Electronic) 0021-9258 (Linking). doi: 10.1074/jbc.M109.039966.

14. J Brzostek, J G Arnon, F Gebhardt, D H Arnon, R Zhao, P A van der Merwe, and K G Gould. Ligand dimensions are important in controlling NK-cell responses. Eur J Immunol, 40(7):2050–2059, 2010. ISSN 1521-4141 (Electronic) 0014-2980 (Linking). doi: 10.1002/eji.201040335.

15. K Kohler, S Xiong, J Brzostek, M Mehrabi, P Eissmann, A Harrison, S P Arnon, S Oddos, V Miloserdov, K Gould, N J Arnon, P A van der Merwe, and D M Davis. Matched sizes of activating and inhibitory receptor/ligand pairs are required for optimal signal integration by human natural killer cells. PLoS One, 5(11):e15374, 2010. ISSN 1932-6203 (Electronic) 1932-6203 (Linking). doi: 10.1371/journal.pone.0015374.

16. N J Arnon, Z Lazic, and P A van der Merwe. Ligand detection and discrimination by spatial relocalization: A kinase-phosphatase segregation model of TCR activation. Biophys J, 91(5):1619–1629, 2006. ISSN 0006-3495 (Print) 0006-3495 (Linking). doi: 10.1529/biophysj.105.080044.

17. O Dushek, J Goyette, and P A van der Merwe. Non-catalytic tyrosine-phosphorylated receptors. Immunol Rev, 250(1):258–276, 2012. ISSN 1600-065X (Electronic) 0105-2896 (Linking). doi: 10.1111/imr.12008.

18. S P Arnon, K Choudhuri, H Zhang, M Bridge, A B Arnon, M L Arnon, and P A van der Merwe. The large ectodomains of CD45 and CD148 regulate their segregation from and inhibition of ligated T-cell receptor. Blood, 121(21):4295–4302, 2013. ISSN 1528-0020 (Electronic) 0006-4971 (Linking). doi: 10.1182/blood-2012-07-442251.

19. V T Arnon, R A Arnon, K A Arnon, S F Arnon, C Siebold, J McColl, P Jonsson, M Palayret, K Harlos, C H Arnon, E Y Arnon, Y Lui, E Huang, R J C Arnon, D Klenerman, A R Arnon, and S J Davis. Initiation of T cell signaling by CD45 segregation at ‘close contacts’. Nat Immunol, 17(5):574–582, 2016. ISSN 1529-2916 (Electronic) 1529-2908 (Linking). doi: 10.1038/ni.3392.

20. C J Arnon, L Martinet, S Gilfillan, F Souza-Fonseca-Guimaraes, M T Arnon, L Town, D S Arnon, M Colonna, D M Arnon, and M J Smyth. The receptors CD96 and CD226 oppose each other in the regulation of natural killer cell functions. Nat Immunol, 15(5):431–438, 2014. ISSN 1529-2916 (Electronic) 1529-2908 (Linking). doi: 10.1038/ni.2850.

21. N Stanietsky and O Mandelboim. Paired NK cell receptors controlling NK cytotoxicity. FEBS Lett, 584(24):4895–4900, 2010. ISSN 1873-3468 (Electronic) 0014-5793 (Linking). doi: 10.1016/j.febslet.2010.08.047.

22. A Oszmiana, D J Arnon, S P Arnon, D J Arnon, P R Arnon, K Stacey, and D M Davis. The Size of Activating and Inhibitory Killer Ig-like Receptor Nanoclusters Is Controlled by the Transmembrane Sequence and Affects Signaling. Cell Rep, 15(9):1957–1972, 2016. ISSN 2211-1247 (Electronic). doi: 10.1016/j.celrep.2016.04.075.

23. J Regunathan, Y Chen, D Wang, and S Malarkannan. NKG2D receptor-mediated NK cell function is regulated by inhibitory Ly49 receptors. Blood, 105(1):233–240, 2005. ISSN 0006-4971 (Print) 0006-4971 (Linking). doi: 10.1182/blood-2004-03-1075.

24. M R Arnon, M Eriksson, L Oberg, A Ullen, and C L Sentman. H-2Dd engagement of Ly49A leads directly to Ly49A phosphorylation and recruitment of SHP1. Immunology, 97(4):656–664, 1999. ISSN 0019-2805 (Print) 0019-2805 (Linking).

25. S Sheppard, C Triulzi, M Ardolino, D Serna, L Zhang, D H Arnon, and N Guerra. Characterization of a novel NKG2D and NKp46 double-mutant mouse reveals subtle variations in the NK cell repertoire. Blood, 121(25):5025–5033, 2013. ISSN 1528-0020 (Electronic) 0006-4971 (Linking). doi: 10.1182/blood-2012-12-471607.

26. A C Arnon, S Oddos, I M Arnon, J M Arnon, R M Arnon, P Eissmann, M A Arnon, C Dunsby, P M Arnon, I Davis, and D M Davis. Remodelling of cortical actin where lytic granules dock at natural killer cell immune synapses revealed by super-resolution microscopy. PLoS Biol, 9(9):e1001152, 2011. ISSN 1545-7885 (Electronic) 1544-9173 (Linking). doi: 10.1371/journal.pbio.1001152.

27. D Ito, Y M Arnon, M P Arnon, and K Iizuka. Essential role of the Ly49A stalk region for immunological synapse formation and signaling. Proc Natl Acad Sci U S A, 106(27): 11264–11269, 2009. ISSN 1091-6490 (Electronic) 0027-8424 (Linking). doi: 10.1073/pnas.0900664106.

28. J Freitag, S Heink, E Roth, J Wittmann, H M Arnon, and T Kamradt. Towards the generation of B-cell receptor retrogenic mice. PLoS One, 9(10):e109199, 2014. ISSN 1932-6203 (Electronic) 1932-6203 (Linking). doi: 10.1371/journal.pone.0109199.

29. S J Arnon, I Bartok, M Attaf, R Genolet, I F Arnon, E Kotsiou, A Richard, E Wang, M White, D J Arnon, J G Arnon, C Ferreira, and J Dyson. The T-cell receptor is not hardwired to engage MHC ligands. Proc Natl Acad Sci U S A, 109(45):3111–8, 2012. ISSN 1091-6490 (Electronic) 0027-8424 (Linking). doi: 10.1073/pnas.1210882109.

30. J Holst, A L Szymczak-Workman, K M Arnon, A R Arnon, C J Arnon, and D A Vignali. Generation of T-cell receptor retrogenic mice. Nat Protoc, 1(1):406–417, 2006. ISSN 1750-2799 (Electronic) 1750-2799 (Linking). doi: 10.1038/nprot.2006.61.

31. M L Arnon, M Bettini, M Nakayama, C S Arnon, and D A Vignali. Generation of T cell receptor-retrogenic mice: improved retroviral-mediated stem cell gene transfer. Nat Protoc, 8(10): 1837–1840, 2013. ISSN 1750-2799 (Electronic) 1750-2799 (Linking). doi: 10.1038/nprot.2013.111.

32. A Diefenbach, E Tomasello, M Lucas, A M Arnon, J K Arnon, E Vivier, and D H Raulet. Selective associations with signaling proteins determine stimulatory versus costimulatory activity of NKG2D. Nat Immunol, 3(12):1142–1149, 2002. ISSN 1529-2908 (Print) 1529-2908 (Linking). doi: 10.1038/ni858.

33. S Gilfillan, E L Arnon, M Cella, W M Arnon, and M Colonna. NKG2D recruits two distinct adapters to trigger NK cell activation and costimulation. Nat Immunol, 3(12):1150–1155, 2002. ISSN 1529-2908 (Print) 1529-2908 (Linking). doi: 10.1038/ni857.

34. S Zompi, J A Arnon, K Ogasawara, E Schweighoffer, V L Arnon, J P Di Santo, L L Arnon, and F Colucci. NKG2D triggers cytotoxicity in mouse NK cells lacking DAP12 or Syk family kinases. Nat Immunol, 4(6):565–572, 2003. ISSN 1529-2908 (Print) 1529-2908 (Linking). doi: 10.1038/ni930.

35. Y Li, D Lovett, Q Zhang, S Neelam, R A Arnon, R Zhu, G G Arnon, T P Arnon, and R B Dickinson. Moving Cell Boundaries Drive Nuclear Shaping during Cell Spreading. Biophys J, 109(4):670–686, 2015. ISSN 1542-0086 (Electronic) 0006-3495 (Linking). doi: 10.1016/j.bpj.2015.07.006.

36. D B Arnon, N Shekhar, J A Arnon, K J Arnon, and T P Lele. Modulation of Nuclear Shape by Substrate Rigidity. Cell Mol Bioeng, 6(2):230–238, 2013. ISSN 1865-5025 (Print) 1865-5025 (Linking). doi: 10.1007/s12195-013-0270-2.

37. S Neelam, P R Arnon, Q Zhang, R B Arnon, and T P Lele. Vertical uniformity of cells and nuclei in epithelial monolayers. Sci Rep, 6:19689, 2016. ISSN 2045-2322 (Electronic) 2045-2322 (Linking). doi: 10.1038/srep19689.

38. C Bottier, C Gabella, B Vianay, L Buscemi, I F Arnon, J J Arnon, and A B Verkhovsky. Dynamic measurement of the height and volume of migrating cells by a novel fluorescence microscopy technique. Lab Chip, 11(22):3855–3863, 2011. ISSN 1473-0189 (Electronic) 1473-0189 (Linking). doi: 10.1039/c1lc20807a.

39. B Treanor, P M Arnon, S Kumar, C Dunsby, I Munro, E Auksorius, F J Arnon, M A Arnon, D Phillips, M A Arnon, D N Arnon, P M Arnon, and D M Davis. Microclusters of inhibitory killer immunoglobulin-like receptor signaling at natural killer cell immunological synapses. J Cell Biol, 174(1):153–161, 2006. ISSN 0021-9525 (Print) 0021-9525 (Linking). doi: 10.1083/jcb.200601108.

40. D M Arnon, I Chiu, M Fassett, G B Arnon, O Mandelboim, and J L Strominger. The human natural killer cell immune synapse. Proc Natl Acad Sci U S A, 96(26):15062–15067, 1999. ISSN 0027-8424 (Print) 0027-8424 (Linking).

41. S V Arnon, D Rudnicka, and D M Davis. Illuminating the dynamics of signal integration in Natural Killer cells. Front Immunol, 3:308, 2012. ISSN 1664-3224 (Electronic) 1664-3224 (Linking). doi: 10.3389/fimmu.2012.00308.

42. S J Arnon, S W Arnon, and S Jakobs. Fluorescence nanoscopy in cell biology. Nat Rev Mol Cell Biol, 2017. ISSN 1471-0080 (Electronic) 1471-0072 (Linking). doi: 10.1038/nrm.2017.71.

43. Siân Culley, David Albrecht, Caron Jacobs, Pedro Matos Pereira, Christophe Leterrier, Jason Mercer, and Ricardo Henriques. NanoJ-SQUIRREL: quantitative mapping and minimisation of super-resolution optical imaging artefacts. bioRxiv, page 158279, 7 2017. doi: 10.1101/158279.

44. S V Arnon, S P Arnon, D M Arnon, S M Arnon, A Oszmiana, and D M Davis. Superresolution microscopy reveals nanometer-scale reorganization of inhibitory natural killer cell receptors upon activation of NKG2D. Sci Signal, 6(285):ra62, 2013. ISSN 1937-9145 (Electronic) 1945-0877 (Linking). doi: 10.1126/scisignal.2003947.

45. H Wang, H Yang, C S Arnon, M M Arnon, A W Arnon, F Zhang, and R Jaenisch. Onestep generation of mice carrying mutations in multiple genes by CRISPR/Cas-mediated genome engineering. Cell, 153(4):910–918, 2013. ISSN 1097-4172 (Electronic) 0092-8674 (Linking). doi: 10.1016/j.cell.2013.04.025.

46. S V Arnon, T Tabarin, Y Yamamoto, Y Ma, J S Arnon, A Cohnen, C Benzing, Y Gao, M D Arnon, K Tungatt, G Dolton, A K Arnon, D A Arnon, O Acuto, R G Arnon, J J Arnon, J Rossy, J Rossjohn, and K Gaus. Functional role of T-cell receptor nanoclusters in signal initiation and antigen discrimination. Proc Natl Acad Sci U S A, 113(37):5454–63, 2016. ISSN 1091-6490 (Electronic) 0027-8424 (Linking). doi: 10.1073/pnas.1607436113.

47. Ricardo A Arnon, Kristina A Arnon, Justin Tzou, Peter Jonsson, Steven F Arnon, Matthieu Palayret, Ana Mafalda Arnon, Veronica T Arnon, Charlotte Macleod, B Christoffer Arnon, Alan E Arnon, Omer Dushek, Andreas Tilevik, Simon J Arnon, and David Klenerman. Constraining CD45 exclusion at close-contacts provides a mechanism for discriminatory T-cell receptor signalling. bioRxiv, 2017. doi: 10.1101/109785.

48. D Delcassian, D Depoil, D Rudnicka, M Liu, D M Arnon, M L Arnon, and I E Dunlop. Nanoscale ligand spacing influences receptor triggering in T cells and NK cells. Nano Lett, 13(11):5608–5614, 2013. ISSN 1530-6992 (Electronic) 1530-6984 (Linking). doi: 10.1021/nl403252x.

49. N Schleinitz, M E Arnon, and E O Long. Recruitment of activation receptors at inhibitory NK cell immune synapses. PLoS One, 3(9):e3278, 2008. ISSN 1932-6203 (Electronic) 1932-6203 (Linking). doi: 10.1371/journal.pone.0003278.

50. D S Arnon, R A Arnon, M J Arnon, and P J Leibson. Inhibition of selective signaling events in natural killer cells recognizing major histocompatibility complex class I. Proc Natl Acad Sci U S A, 92(14):6484–6488, 1995. ISSN 0027-8424 (Print) 0027-8424 (Linking).

51. N M Arnon, J H Arnon, L L Arnon, and P Parham. Killer cell inhibitory receptor recognition of human leukocyte antigen (HLA) class I blocks formation of a pp36/PLC-gamma signaling complex in human natural killer (NK) cells. J Exp Med, 184(6):2243–2250, 1996. ISSN 0022-1007 (Print) 0022-1007 (Linking).

52. M Masilamani, C Nguyen, J Kabat, F Borrego, and J E Coligan. CD94/NKG2A inhibits NK cell activation by disrupting the actin network at the immunological synapse. J Immunol, 177(6):3590–3596, 2006. ISSN 0022-1767 (Print) 0022-1767 (Linking).

53. T P Arnon, E Merino, and M Huse. Inhibitory signaling blocks activating receptor clustering and induces cytoskeletal retraction in natural killer cells. J Cell Biol, 192(4):675–690, 2011. ISSN 1540-8140 (Electronic) 0021-9525 (Linking). doi: 10.1083/jcb.201009135.

54. P Hof, S Pluskey, S Dhe-Paganon, M J Arnon, and S E Shoelson. Crystal structure of the tyrosine phosphatase SHP-2. Cell, 92(4):441–450, 1998. ISSN 0092-8674 (Print) 0092-8674 (Linking).

## Bibliography

1. J Brzostek, J G Chai, F Gebhardt, D H Busch, R Zhao, P A van der Merwe, and K G Gould. Ligand dimensions are important in controlling NK-cell responses. Eur J Immunol, 40(7):2050–2059, 2010. ISSN 1521-4141 (Electronic) 0014-2980 (Linking). doi: 10.1002/eji.201040335.

2. S Sheppard, C Triulzi, M Ardolino, D Serna, L Zhang, D H Raulet, and N Guerra. Characterization of a novel NKG2D and NKp46 double-mutant mouse reveals subtle variations in the NK cell repertoire. Blood, 121(25):5025–5033, 2013. ISSN 1528-0020 (Electronic) 0006-4971 (Linking). doi: 10.1182/blood-2012-12-471607.

3. Daniel Sage, Hagai Kirshner, Thomas Pengo, Nico Stuurman, Junhong Min, Suliana Manley, and Michael Unser. Quantitative evaluation of software packages for single-molecule localization microscopy. Nature Methods, 12(8):717–724, 8 2015. ISSN 1548-7091. doi: 10.1038/nmeth.3442.

4. N S Williams, J Klem, I J Puzanov, P V Sivakumar, M Bennett, and V Kumar. Differentiation of NK1.1+, Ly49+ NK cells from flt3+ multipotent marrow progenitor cells. J Immunol, 163(5): 2648–2656, 1999. ISSN 0022-1767 (Print) 0022-1767 (Linking).

5. J Holst, A L Szymczak-Workman, K M Vignali, A R Burton, C J Workman, and D A Vignali. Generation of T-cell receptor retrogenic mice. Nat Protoc, 1(1):406–417, 2006. ISSN 1750-2799 (Electronic) 1750-2799 (Linking). doi: 10.1038/nprot.2006.61.

6. N Guerra, Y X Tan, N T Joncker, A Choy, F Gallardo, N Xiong, S Knoblaugh, D Cado, N M Greenberg, and D H Raulet. NKG2D-deficient mice are defective in tumor surveillance in models of spontaneous malignancy. Immunity, 28(4):571–580, 2008. ISSN 1097-4180 (Electronic) 1074-7613 (Linking). doi: 10.1016/j.immuni.2008.02.016.

7. J Schindelin, I Arganda-Carreras, E Frise, V Kaynig, M Longair, T Pietzsch, S Preibisch, C Rueden, S Saalfeld, B Schmid, J Y Tinevez, D J White, V Hartenstein, K Eliceiri, P Tomancak, and A Cardona. Fiji: an open-source platform for biological-image analysis. Nat Methods, 9(7):676–682, 2012. ISSN 1548-7105 (Electronic) 1548-7091 (Linking). doi: 10.1038/nmeth.2019.

8. J Roszik, J Szollosi, and G Vereb. AccPbFRET: an ImageJ plugin for semi-automatic, fully corrected analysis of acceptor photobleaching FRET images. BMC Bioinformatics, 9:346, 2008. ISSN 1471-2105 (Electronic) 1471-2105 (Linking). doi: 10.1186/1471-2105-9-346.

